# Bacterial transcriptional repressor NrdR – a flexible multifactorial nucleotide sensor

**DOI:** 10.1101/2024.09.04.609659

**Authors:** Inna Rozman Grinberg, Ornella Bimaï, Saher Shahid, Lena Philipp, Markel Martínez-Carranza, Ipsita Banerjee, Daniel Lundin, Pål Stenmark, Britt-Marie Sjöberg, Derek T. Logan

## Abstract

NrdR is a bacterial transcriptional repressor consisting of a Zn-ribbon domain followed by an ATP-cone domain. Understanding its mechanism of action could aid the design of novel antibacterials. NrdR binds specifically to two “NrdR boxes” upstream of ribonucleotide reductase operons, of which *Escherichia coli* has three: nrdHIEF, nrdDG and nrdAB, where we identified a new box. We show that *E. coli* NrdR (EcoNrdR) has similar binding strength to all three sites when loaded with ATP plus dATP or equivalent diphosphate combinations. No other combination of nucleotides promotes binding to DNA. We present crystal structures of EcoNrdR-ATP-dATP and EcoNrdR-ADP-dATP, which are the first high-resolution crystal structures of an NrdR. We have also determined cryo-EM structures of DNA-bound EcoNrdR-ATP-dATP and novel filaments of EcoNrdR-ATP. Tetrameric forms of EcoNrdR involve alternating interactions between pairs of Zn-ribbon domains and ATP-cones. The structures reveal considerable flexibility in relative orientation of ATP-cones vs Zn-ribbon domains. The structure of DNA-bound EcoNrdR-ATP-dATP shows that significant conformational rearrangements between ATP-cones and Zn-ribbons accompany DNA binding while the ATP-cones retain the same relative orientation. In contrast, ATP-loaded EcoNrdR filaments show rearrangements of the ATP-cone pairs and sequester the DNA-binding residues of NrdR such that they are unable to bind to DNA. Our results, in combination with a previous structural and biochemical study, point to highly flexible EcoNrdR structures that when loaded with the correct nucleotides adapt to an optimal promoter binding conformation.

## Introduction

NrdR is a transcription factor that regulates expression of ribonucleotide reductase (RNR), an essential enzyme that provides DNA building blocks in all living cells (1). NrdR is abundant in bacteria, rare in archaea and lacking in eukaryotes. The NrdR protein consists of an N-terminal Zn-ribbon domain (2) followed by an ATP-cone domain (3), and a C-terminal part of variable length and unknown function. The Zn-ribbon domain of approximately 45 residues consists of two stretches of -CXXC-residues that coordinate the zinc ion. NrdR binds to two highly conserved palindromic 16-bp sequences called NrdR boxes located in the operons of RNR genes. DNA binding is controlled via loading of adenosine nucleotides to the approximately 100-residue ATP-cone domain. Interestingly, an ATP-cone also is well-known as an allosteric domain in many RNRs where it controls the activity of the enzyme by binding ATP or dATP (4,5). Regulation of RNR expression is important for DNA replication and repair, as RNR catalyzes the sole de novo pathway for production of deoxyribonucleotides (dNTPs) (6). To our knowledge this is the first encounter of a highly specific repressor with a horizontally transferable domain that controls both the expression of an enzyme and its catalytic activity.

Three evolutionarily related classes of RNR are known: an oxygen-requiring class I RNR encoding a catalytic subunit and a radical-generating subunit, an adenosylcobalamin-dependent class II RNR, and an oxygen-sensitive class III RNR (7–9). Whereas eukaryotes, with few exceptions, encode only class I RNRs, both bacteria and archaea can encode any of the three classes and in many cases more than one, making transcriptional control of RNR genes necessary both from environmental and metabolic perspectives. *Escherichia coli* NrdR (EcoNrdR), the focus of this study, controls three RNR operons: two class I operons encoding RNRs with different metal requirements and one anaerobically required class III operon (10).

NrdR studies have primarily been focused on the proteins from *Streptomyces coelicolor*, *Escherichia coli* and *Pseudomonas aeruginosa* (1,2,11–14). The NrdR boxes overlap with the RNR polymerase binding site and several studies have shown that deletion of *nrdR* results in a general increase of *nrd* expression (10,14–16). Recently, we solved cryo-EM structures of DNA-bound and unbound *S. coelicolor* NrdR (ScoNrdR) (1). In combination with extensive biochemical studies our results suggested a mechanism of action for NrdR, where an ATP-loaded NrdR forms a dodecamer that inhibits binding to DNA, whereas loading the other nucleotide binding site on NrdR with dATP promotes formation of a tetramer that can bind to DNA. Our high resolution cryo-EM structure of ScoNrdR bound to DNA shows that the tetramer bends DNA in a sharp kink (1).

Here we have used a variety of biochemical studies to delineate the extensive nucleotide binding capacity of *E. coli* NrdR, and to demonstrate which nucleotides are required for its binding to DNA. We also present three high resolution X-ray structures of EcoNrdR loaded with different nucleotides and lower-resolution cryo-EM structures of EcoNrdR bound to DNA and in a filamentous form induced by ATP alone. Our results lead to a hypothesis for the mechanism of action of EcoNrdR that differs from earlier suggested ones (10,13). The evolutionary relationship between RNR regulation at the protein and gene levels through the common ATP-cone is a unique example of a multifactorial nucleotide-binding sensor that can be exploited in development of novel antibiotics.

## Results

### Binding of NrdR to RNR promoter regions

Of the three RNR operons in *E. coli (nrdAB, nrdHIEF, nrdDG),* the two latter encode two characteristic NrdR boxes each (box 1 and box 2) in the promoter regions, with a distance of 15 bp between the boxes (Fig. 1). The number of NrdR boxes in the *nrdAB* promoter region is less obvious. Earlier literature suggested two boxes with a linker distance of 16 bp and overlapping the RNA polymerase binding site. However, earlier experimental results suggested that the upstream box (here called box 0) had less influence on binding of NrdR than the downstream box (here called box 1) (10). Interestingly, a potential third NrdR box (here called box 2) exists downstream of the promoter region of *nrdAB* at a distance of 15 bp to the upstream NrdR box, forming an alternative NrdR box pair. Whereas all other NrdR boxes in *E. coli* have a central - AT-sequence, the new *nrdAB* box 2 has a non-canonical central - AG-sequence (Fig. 1).

**Figure 1.**
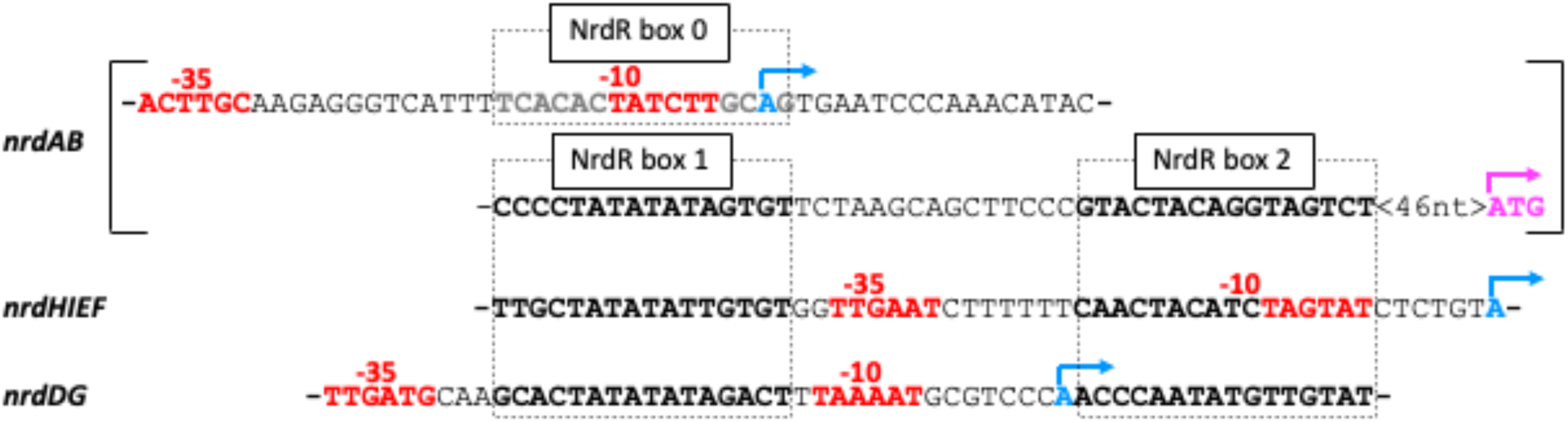
Promoter regions of RNR genes in *E. coli*. NrdR box regions are in bold and boxed, RNA polymerase interaction regions −10 and −35 are in red, transcription start sites in blue with arrows showing transcription direction, and translation start site in violet with arrow. A previously denoted NrdR box (here called box 0) in the −10 region of the *nrdAB* promoter is indicated in grey.

To determine the binding affinity of NrdR to each of the promoter regions, we first established which nucleotide effectors are necessary for binding using MST. As was earlier shown for *S. coelicolor* NrdR (1), combinations of dATP with either ATP or ADP were required for specific binding of *E. coli* NrdR to 57-bp DNA fragments of the promoter regions of *nrdDG* and *nrdHIEF* (Table 1). NrdR was unable to bind to the negative control *ribX* promoter, which lacks NrdR boxes (Supplementary Figure S4).

**Table 1.** Promoter binding constants for effector nucleotide-loaded *E. coli* NrdR.

| Effector | *nrdHIEF* promoter<br>$K_D^a$ [nM] | *nrdDG* promoter<br>$K_D^a$ [nM] | *nrdAB* promoter<br>box 1+2<br>$K_D^a$ [nM] | *nrdAB* promoter<br>box 0+1<br>$K_D^a$ [nM] |
| --- | --- | --- | --- | --- |
| dATP + ATP | 138 ± 32 | 160 ± 37 | 113 ± 27 | > 6 000 |
| dATP + ADP | 187 ± 36 | 152 ± 42 | 96 ± 18 | > 6 000 |
| dATP + AMP | > 5 000 | > 5 000 | > 10 000 | > 10 000 |
| ATP | n.d.<sup>b</sup> | n.d.<sup>b</sup> | n.d.<sup>b</sup> | n.d.<sup>b</sup> |
| ADP | > 100 000 | n.d.<sup>b</sup> | > 5 000 | n.d.<sup>b</sup> |

<sup>a</sup>Determined using MST as described in Materials and Methods. $K_D$ and standard deviation were calculated using fits from at least three individual titrations.
<sup>b</sup>n.d.; not detected

Binding to the originally predicted NrdR boxes in the *nrdAB* promoter region (boxes 0 and 1) was very weak (40 to 50 times lower affinity than for the *nrdDG* and *nrdEF* promoter regions). No binding to this probe was detected even in the presence of other nucleotides or their combinations (Supplementary Table S1). We then tested binding of NrdR to the alternative NrdR box motif (box 1 and box 2) in the *nrdAB* promoter and detected high affinity binding of NrdR (Table 1; Supplementary Fig. S4). Our results are consistent with earlier results from electrophoretic mobility shift assays using a 185 bp *nrdAB* promoter region containing all three NrdR boxes (box 0, box 1, box 2) showing that mutation of box 0 did not affect the binding of NrdR, whereas mutation of box 1 abolished binding (10). We therefore consider NrdR boxes 1 and 2 in the *nrdAB* downstream promoter region to constitute the true NrdR binding site (Fig. 1).

As previously reported (2,13), as-purified NrdR contained ∼0.5 mol nucleotide/ mol NrdR, primarily ADP and AMP (Supplementary Fig. S5). To explore which nucleotides promote NrdR binding to DNA we used apo-protein from which nucleotides that co-purified with the protein have been removed. When single nucleotides (ATP, dATP, ADP, dADP, AMP or dAMP) were added to the reaction mixture, NrdR did not bind DNA, or in rare cases bound with significantly lower affinity (Table 1, Supplementary Table S1, Supplementary Figure S6). After establishing that a combination of two nucleotides is required for binding, we tested the effects of all possible adenosine mono-, di- and triphosphate combinations on NrdR binding to each of the promoters (Supplementary Table S1, Supplementary Figure S7). While most combinations resulted in no binding or binding with very low affinity, the combination of dADP and ADP to NrdR resulted in approximately similar DNA affinities as the combination of dATP with either ATP or ADP. While dATP/ADP-loaded NrdR is likely to exist *in vivo* at low cellular energy levels (Supplementary Figure S5), the physiological relevance of the combination dADP/ADP is questionable, since the intracellular concentration of dNDP in general is very low (30,31). The addition of cyclic di-AMP alone or in combination with ADP as well as addition of cAMP did not promote NrdR binding to DNA (Supplementary Table S1, Supplementary Figure S8).

### Oligomeric states of NrdR in the presence of allosteric effectors

To elucidate the oligomeric states of *E. coli* NrdR, size exclusion chromatography (SEC) was used. In the absence of nucleotide effectors, NrdR eluted as a dimer (Fig. 2). In the presence of the nucleotide combinations dATP plus ATP or dATP plus ADP, which promote DNA binding, NrdR eluted mainly as a tetramer. ADP-loaded NrdR also eluted as a tetramer, as did NrdR in the presence of the combination of AMP plus dAMP. ATP-loaded NrdR, a form unable to bind DNA, eluted as a dodecamer at low concentrations of magnesium chloride (0.1 - 0.2 mM), but in the void volume of the Superdex 200 column when the magnesium concentration was increased (≥1 mM; Fig. 2). Increasing concentrations of ATP-loaded NrdR resulted in increasing amounts eluting in the void volume (Supplementary Figure S9). Also dATP/ATP-loaded NrdR and dATP/ADP-loaded NrdR showed increasing molecular mass complexes at increasing protein concentration, but remained soluble at concentrations used for crystallography (∼550 μM), which produced crystals of tetramers (see below). (Supplementary Figure S9).

**Figure 2.**
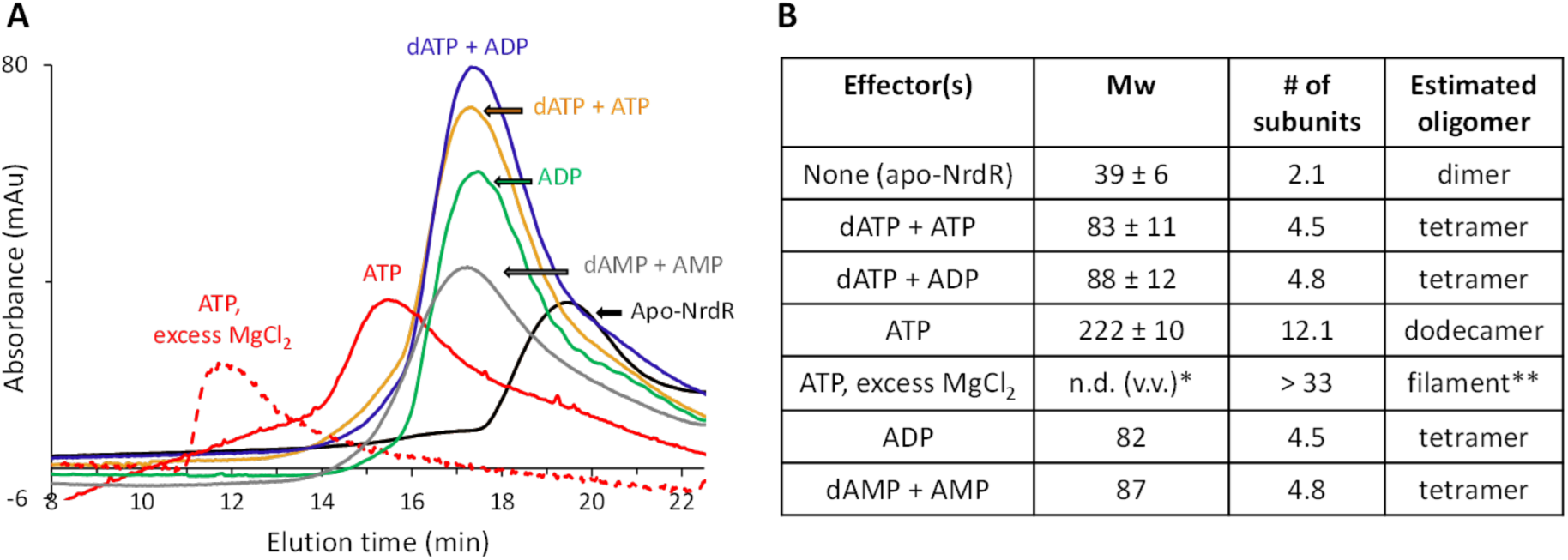
Size exclusion chromatography of *E. coli* NrdR in the presence or absence of nucleotide effectors. **(A)** Representative chromatograms are shown, in which 50 μl and 93 μl of 55 μM apo-NrdR pre-incubated with ATP and with other effectors respectively, were loaded on Superdex 200 PC 3.2/30 (with a total volume of 2.4 ml) connected to a Shimadzu HPLC system. **(B)** Molecular mass of the eluted protein, calculated based on protein standards, the number of monomeric subunits in the eluted complex and the estimated oligomeric state of NrdR. NrdR was pre-incubated with the indicated nucleotides (or without nucleotides) for 10-60 min. at 8 °C and the samples were applied to SEC as described in Materials and Methods. Standard deviations were determined using data obtained from a minimum of three experimental runs. * n.d. (v.v.): not determined, because the sample elutes in the void volume of the column. ** based on the cryo EM structure.

### NrdR crystal structures

Structures of *E. coli* NrdR were solved in three co-crystallised nucleotide-loaded states: with dATP and the non-hydrolysable ATP analogue AMPPNP, with dATP and ADP and with dATP and ATP that hydrolyzed to ADP during crystallisation. Data and structure quality statistics are listed in Supplementary Table S2. All of the structures confirm a two-domain arrangement consisting of an N-terminal Zn-ribbon domain (residues 1-44) and a C-terminal ATP-cone domain (50–150) joined by a flexible linker (45-49; Fig. 3). In the Zn-ribbon, the Zn^2+^ ions are coordinated by Cys3, Cys6, Cys31 and Cys34. EcoNrdR contains the same type of ATP-cone domain recently identified in the structure of *S. coelicolor* NrdR (1), with two nucleotide binding sites, an “inner” and an “outer” site, having differing specificities. The flexible linker allows the two domains to adopt multiple different relative conformations.

**Figure 3.**
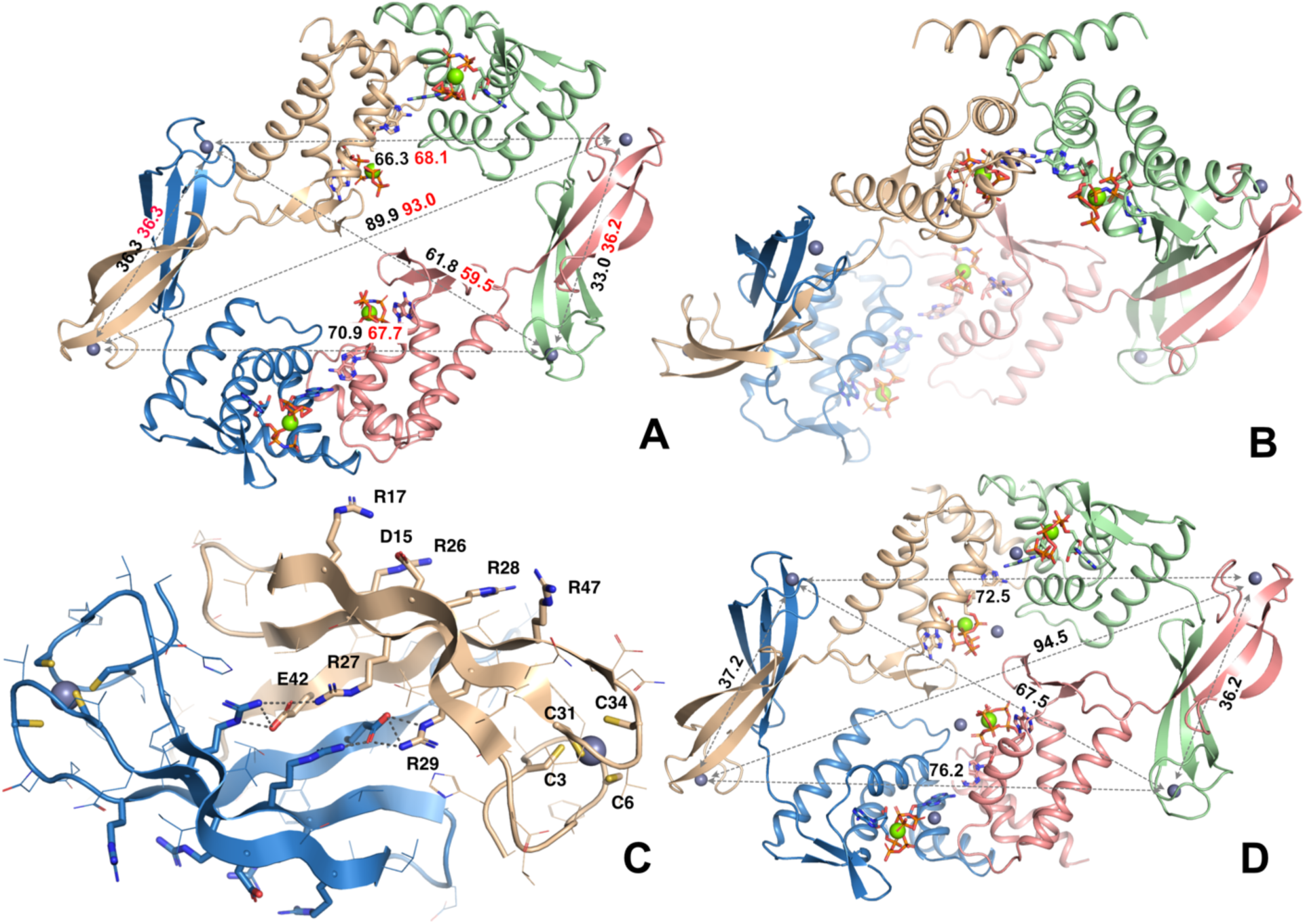
Crystal structures of the AMPPNP/dATP-loaded and ATP-dATP-loaded *E. coli* NrdR tetramers. Chain A is coloured beige, chain B green, chain C pink and chain D blue. Nucleotides are shown as sticks. A) View of the AMPPNP/dATP-loaded tetramer down the 2-fold symmetry axis, from the DNA-binding side. Distances between pairs of Zn^2+^ ions in the DNA-binding Zn-ribbon domains, which help define the dimensions of the tetramer, are shown in black and red text for the two independent tetramers in the asymmetric unit respectively. B) View in which the local 2-fold symmetry axis of the upper pair of ATP-cones is vertical. In this view the distinct orientations of the Zn-ribbon domains relative to the ATP-cones in the blue and pink chains are evident. C) Closeup view of a typical pair of Zn-ribbon domains. All side chains are shown as lines. The cysteine ligands of the Zn^2+^ ion, conserved Arg residues and Asp15 on the DNA-binding surface are shown as sticks. Salt bridges that stabilise the dimer interface are shown as grey dashes and sticks. Zn^2+^ ions are shown as grey spheres. For clarity only one monomer is labelled. D) Overview of the ATP-dATP-loaded EcNrdR tetramer in the same view as panel A). Four additional Zn^2+^ ions bound at the nucleotide sites are shown as grey spheres.

### AMPPNP/dATP-loaded EcoNrdR

In complex with dATP and AMPPNP, EcoNrdR is a tetramer consisting of two tightly intertwined dimers. The crystallographic asymmetric unit contains two tetramers that differ slightly in the relative orientation of the ATP-cone and Zn-ribbon domains, presumably due to crystal packing effects. This further highlights the interdomain flexibility of NrdR. The interactions between the Zn-ribbon domains feature a continuous 6-stranded intermolecular β-sheet where residues 39-45 from the third β-strand make an antiparallel interaction. This is stabilised by symmetrical sets of salt bridges and hydrogen bonds around a pseudo-2-fold axis, between a cluster of charged residues on the surface of the β-sheet, in particular Arg27, Arg29 and Glu42 (Fig. 3C). These interactions, as well as the known tendency of structures with exposed β-strands to aggregate in solution (32), make it reasonable to suppose that the EcoNrdR dimers observed in apo-NrdR are associated through their Zn-ribbons and that tetramers are formed by the interaction of ATP-cones under the influence of adenosine nucleotides. The conformations of the Zn-ribbon pairs are almost identical in all the crystal structures of EcoNrdR (Supplementary Figure S10), reinforcing the hypothesis that these pairs act as rigid structural units.

The AMPPNP-dATP loaded tetramer has overall pseudo-C2 symmetry (Fig. 3A) but viewing the ATP-cone pairs as symmetrical dimers (Fig. 3B), it is clear that the Zn-ribbons have two distinct orientations relative to them in the tetramer. A simple way to classify the relative domain orientations is through the angle subtended by the Cɑ atoms of residues 81 and 98 (that begin and end helix 4 of the ATP-cone domain) and the Zn^2+^ ion in the Zn-ribbon domain (Fig. 4C).

**Figure 4.**
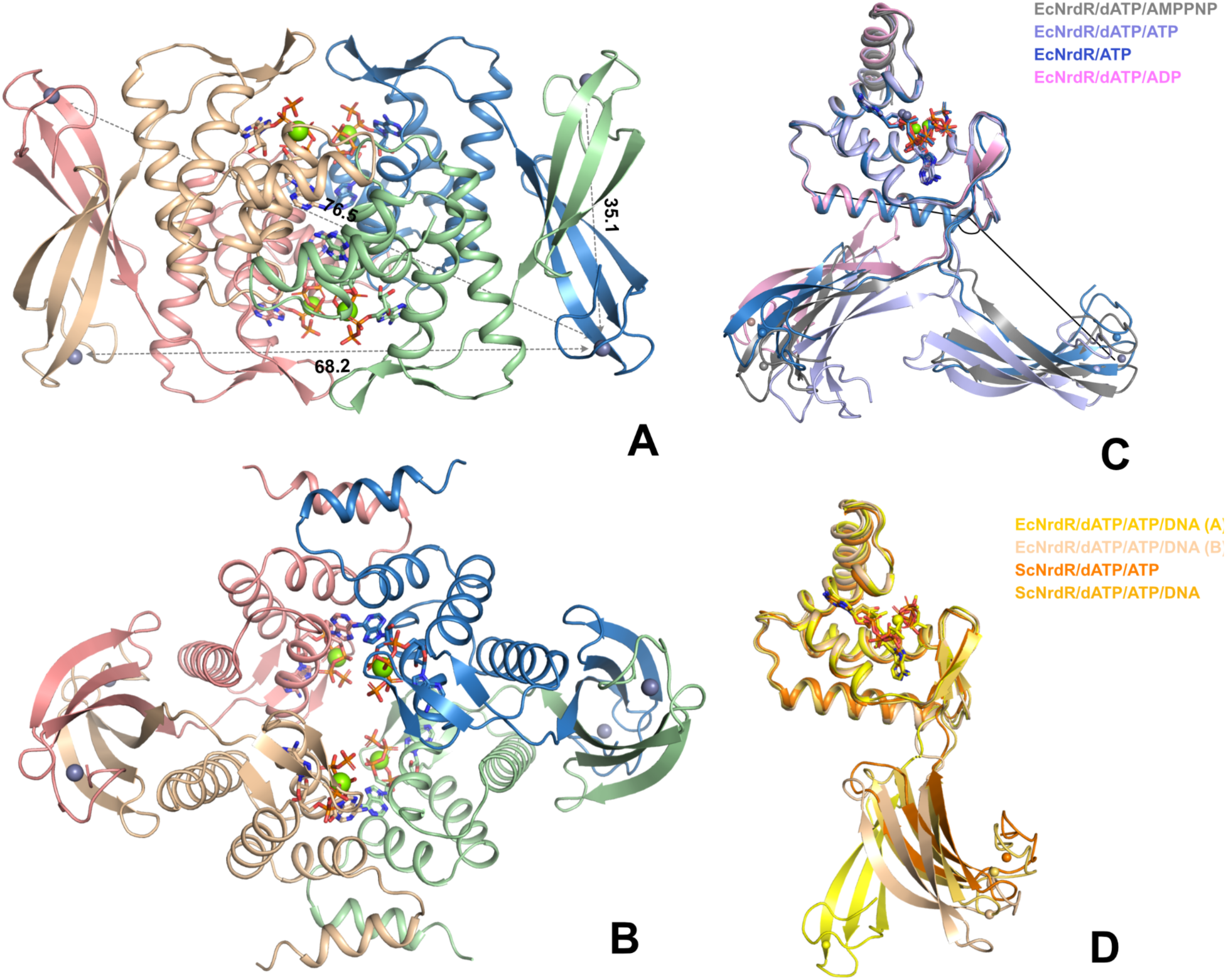
The tetramer of EcoNrdR in its ADP/dATP-bound state, generated by crystallographic symmetry from a single chain in the asymmetric unit. A) and B) show two perpendicular views chosen to be similar to those in Figure 3 A) and B). The molecular representation is also as in Figure 3. Distances between relevant pairs of Zn^2+^ ions in the Zn-ribbon domains are shown. Only one set of distances is shown as the others are equivalent by symmetry. C) Relative orientations of the Zn-ribbon domains relative to the ATP-cone domains in all states of EcoNrdR except the DNA-bound state. The monomers have been superimposed on their ATP-cone domains. Nucleotides are shown as sticks.

One conformation (beige) is fully extended (angle ∼110°) and the other (green) has the Zn ribbon at a ∼45° angle. Thus, the two domains have at least two low-energy states relative to each other. Superposition of the eight chains in the two independent tetramers reveals that the two relative orientations are not unique but rather define clusters (Supplementary Figure S11 and Supplementary Table S3). These lead to the slightly different overall dimensions of the two tetramers (Fig. 3A) and indicate considerable flexibility despite identical nucleotide binding to the ATP-cones and relative orientations of the latter. As will be seen below, further relative orientations exist in other functional states. Interestingly, both relative domain orientations differ distinctly from those seen in the ScNrdR tetramer, which has only slightly different relative dimensions (1). For example, the distances between Zn^2+^ ions in symmetrical pairs of Zn-ribbon dimers in the EcoNrdR tetramer, representing the two binding sites for the NrdR boxes, are on average ∼61 Å and ∼91 Å (Fig. 3A). This compares to 78-79 Å in the dATP/ADP-bound ScNrdR octamer and 82 Å in the DNA-bound tetramer. This suggests that a significant rearrangement of the EcoNrdR tetramer takes place upon DNA binding.

The Zn-ribbon domains are exposed to solvent. The highly conserved residues identified as being essential for DNA recognition in the *S. coelicolor* NrdR structure (1), namely Asp15, Arg17, Arg26, Arg28 and Arg37, are exposed on the outer face of each Zn-ribbon pair in EcoNrdR (Fig. 3C).

### ATP/dATP-loaded SeMet-EcoNrdR

At an early stage in the structure solution, we crystallised Se-Met substituted EcoNrdR in the presence of ATP and dATP. This form crystallised in a different unit cell, with only one tetramer in the asymmetric unit, and the structure was solved at 3.1 Å resolution. Interestingly, this tetramer has a slightly different form than the one loaded with AMPPNP-dATP (Fig. 3D) in which nucleotides in the two halves of the tetramer are in closer proximity than in the AMPPNP-dATP loaded tetramer. The relative orientation of ATP-cone domains is the same, but the Zn ribbon domains have a different relative orientation. Nevertheless, these fall into the two distinct clusters seen in the AMPPNP-dATP-bound tetramers (Supplementary Figure S11). Strong electron density is observed between the α- and γ-phosphate groups of each of the four dATP molecules that can be interpreted as a Zn^2+^ ion from the crystallisation medium (Supplementary Figure S12). Residue Asn55 from the tip of the ꞵ-hairpin of an ATP-cone on one side of the tetramer reaches over to the other side to coordinate the γ-phosphate group of one of the dATP molecules. However, the tetramer is asymmetrical, and this interaction occurs only once.

The SeMet-ATP-dATP structure shows small differences in nucleotide coordination compared to the AMPPNP-dATP complex. The nucleotide pair appears to lack an internal Mg^2+^ ion. This may be a result of binding of Zn^2+^ from the crystallisation solution to the outside of the dATP molecule (Supplementary Figure S12). The γ-phosphate group of ATP partly occupies the space vacated by the Mg^2+^ ion. However, since AMPPNP coordination in EcoNrdR is almost identical to that of ATP in ScoNrdR, we believe that the ATP binding mode seen in this structure may be partially an artefact of the presence of Zn^2+^. Thus, the AMPPNP model, at higher resolution, is valid for analysis of biochemical results involving ATP. Since the Zn^2+^ ion may have affected the tetramer conformation, we limit our interpretation to the observation that the ATP-dATP loaded EcoNrdR tetramer is clearly highly malleable.

### ADP/dATP-loaded EcoNrdR

In initial experiments, NrdR was co-crystallized with dATP and ATP and the structure determined from anisotropic data to a maximum resolution of 2.44 Å. However, the structure revealed that the ATP had been hydrolysed to ADP during crystallisation. The asymmetric unit contains two chains, A and B. At first glance the crystal packing suggests an octameric arrangement, but the octamer is not resolvable into two tetrameric halves like the ScoNrdR octamer. On closer inspection the octamer can be decomposed into compact tetramers generated by crystal symmetry operations from chain B alone, decorated with infinite chains of NrdR monomers (chain A), held together by alternating interactions between pairs of Zn-ribbon domains and pairs of ATP-cone domains (Supplementary Fig. S13). Here we will focus on the structure of the compact tetramer, which is consistent with the SEC experiments that show the same oligomeric state for both dATP-ATP and dATP-ADP.

The compact ADP/dATP-bound tetramer is more symmetrical than the AMPPNP/dATP tetramer, with D2 symmetry (Fig. 4A, 4B). The angles between ATP-cones and Zn-ribbons are thus the same in all monomers. As in all other NrdR structures, the distance between Zn^2+^ ions in a Zn-ribbon pair is ∼35 Å. The distance between Zn^2+^ ions in the two NrdR-box binding sites is 76.5 Å. The ATP-cone pairs are essentially identical to those in the AMPPNP-dATP tetramer, with an rmsd in Cα positions of only 0.3 Å over 152 equivalent residues. Likewise, the nucleotide binding modes are identical, with the exception of the missing γ-phosphate group in ADP (Supplementary Fig. S14). In contrast, the Zn-ribbon domain has a completely different orientation relative to the ATP-cone (Fig. 4C). This compact tetramer is incompatible with DNA binding without major rearrangement, as will be shown below.

Representatives of the two clusters of conformations found in each tetrameric state are shown in the same colour. Chains of type B cluster to the left and chains of type A cluster to the right. The filled black lines indicate the angle subtended by the CA atoms of residues 81 and 98, and the Zn^2+^ ion, which are tabulated for all conformations of EcoNrdR and ScoNrdR in Supplementary Table S3. D) Comparison of the Zn-ribbon orientations in the DNA-bound states of EcoNrdR and ScoNrdR. A much smaller conformational change is observed between the free and DNA-bound forms for ScoNrdR.

### Cryo-EM structure of the ATP-dATP-loaded EcoNrdR-DNA complex

Attempts to obtain a high-resolution cryo-EM structure of the EcoNrdR-dATP-ATP-DNA complex were confounded by the tendency of the particles to adopt a preferential orientation on the cryo-EM grids. By using non-tilted and tilted micrographs, we obtained a map at 4 Å resolution which was locally sharpened using DeepEMhancer (23). Into this map the DNA from the ScNrdR-dATP-ATP-DNA complex (1) and the ATP-cone and Zn-ribbon domains from the EcoNrdR-dATP-ATP tetramer were separately fitted as rigid bodies (Figure 5A, 5B), without further modification. The ATP-cone and Zn-ribbon pairs fit the map well as rigid bodies. The DNA exhibits the same sharp bend at each NrdR box as observed in the ScNrdR complex.

**Figure 5.**
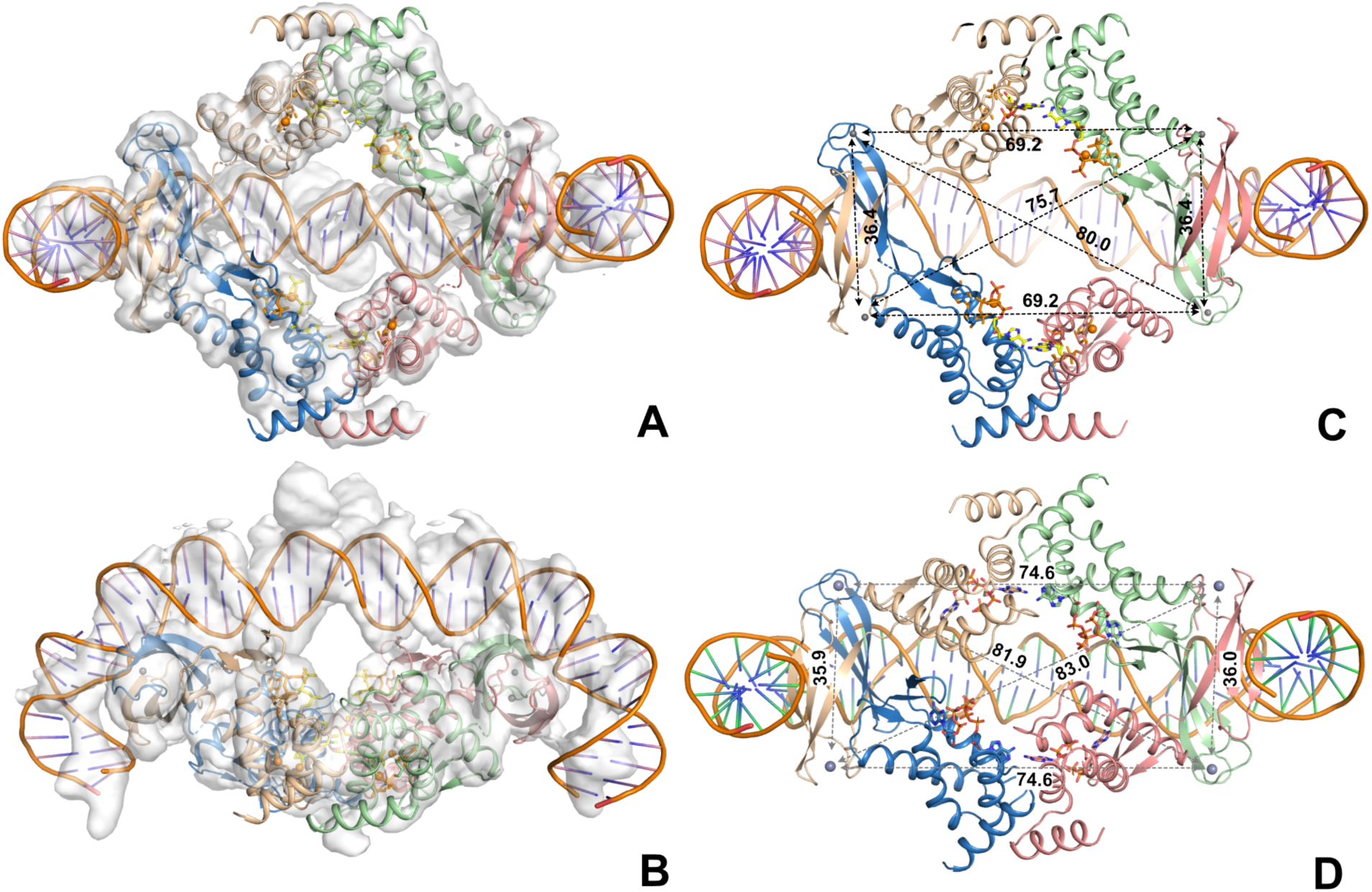
Cryo-EM structure of the EcoNrdR-dATP-ATP-DNA complex. A: View of the complex in the same orientation as Figure 3A. B) View rotated 90° around a horizontal axis. The reason for poor fit of the DNA at the ends of the fragment is not known. C) View of the DNA-bound tetramer without the map. Distances between Zn^2+^ ions that define the overall dimensions of the tetramer are shown. D) The DNA-bound tetramer from *S. coelicolor*.

Interestingly, the relative orientation of the ATP-cones and Zn-ribbons is different to either the EcoNrdR-dATP-ATP tetramer in isolation or to the ScNrdR complex. The maximum diagonal distance between Zn^2+^ ions is reduced from ∼92 Å in the free tetramer to ∼80 Å in the DNA-bound tetramer, a value much closer to that in the *S. coelicor* DNA-bound form (82 Å). However, the ATP-dATP-DNA-bound tetramer of EcoNrdR has a more open form than that of ScNrdR, with a larger cavity in the middle (Figure 5C, 5D, Supplementary Movie S1).

It was previously noted for ScNrdR that the two tetramers that form the two halves of a dATP-ATP-bound octamer and the single tetramer bound to DNA are virtually identical apart from small adjustments of the orientation of ATP-cone and Zn-ribbon domains (1) (see the two orange cartoons in Fig. 4C). In EcoNrdR the situation is very different, and a substantial change in domain orientation occurs upon DNA binding (Supplementary Movie S2). The required conformational change is even larger if starting from the ADP-dATP-bound form (Supplementary Movie S3).

The ADP-dATP-bound tetramer observed in the crystal structure is incompatible with DNA binding without a much larger rearrangement than the one required for binding of the ATP-dATP-bound tetramer. This is consistent with the fact that the relative domain orientation in the ADP-dATP form is most different to any of the others (Fig. 5C). However, this rearrangement is possible without clashing with the DNA.

### Nucleotide binding and specificity in the ATP-cone

The higher resolution of the crystal structures of the AMPPNP/dATP and ADP/dATP-loaded forms of EcoNrdR compared to the cryo-EM structures of ScoNrdR permits detailed insight into the structural features that govern site-specific binding of two nucleotides simultaneously to the ATP-cone. The electron density unambiguously shows that ATP binds to the “inner” site and dATP to the “outer” site (Fig. 6). As in ScoNrdR, the outer-site dATP molecules are found in a head-to-head arrangement across the dimer interface of the pair of ATP-cones. ATP is precluded from binding to the outer site in this state of the ATP-cone domain as its 2’-OH group would clash sterically with the side chain of Phe127 in the last helix of the ATP-cone. A Tyr side chain in the outer site was observed to flip between an “in” and an “out” conformation in the ATP-bound and dATP-bound forms of ScoNrdR respectively (1). The residue is conserved (Tyr131) in EcoNrdR and is, as in the ATP-dATP form of ScoNrdR, in the “out” conformation.

**Figure 6.**
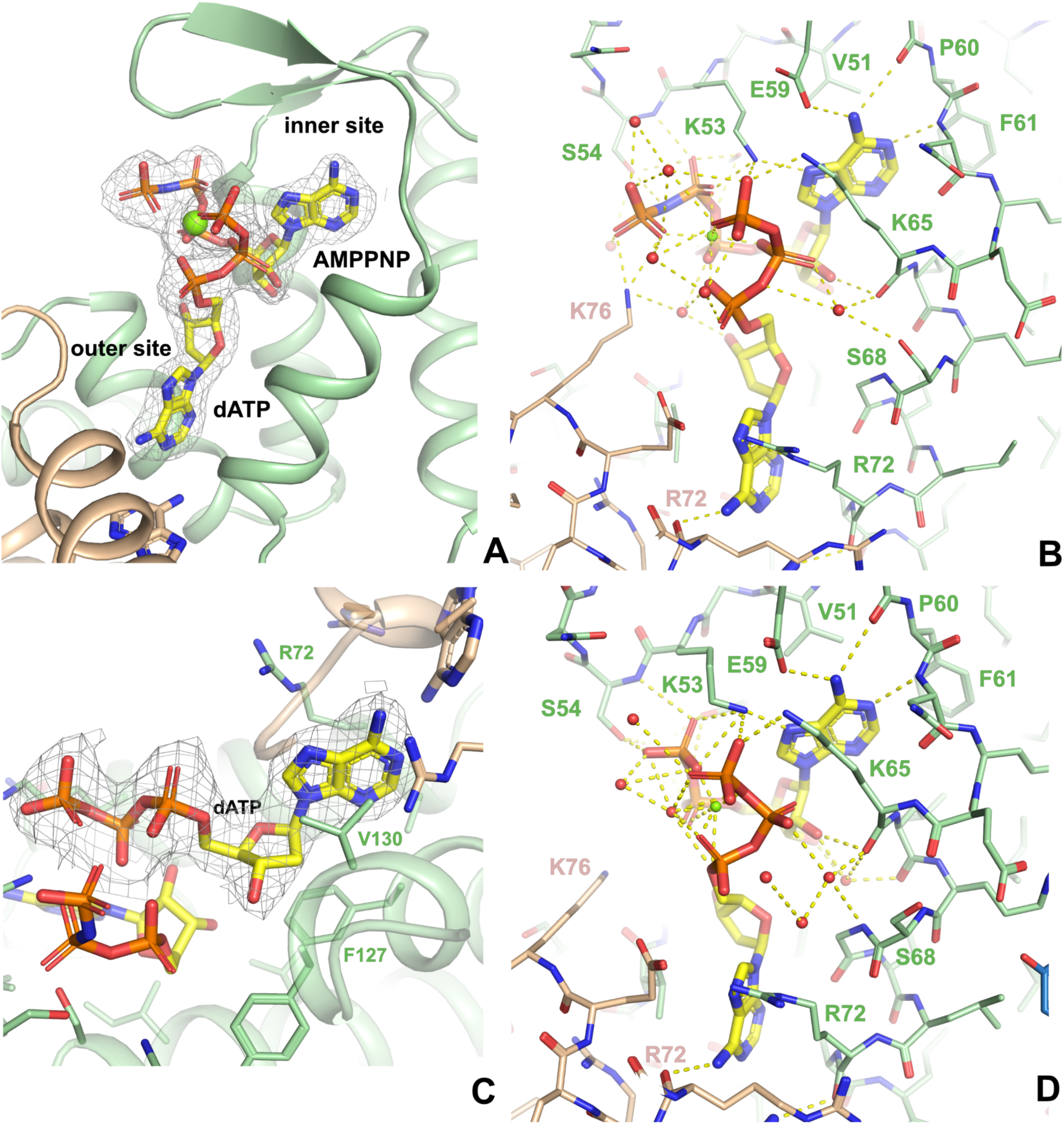
Nucleotide binding in the AMPPNP/dATP complex of EcoNrdR. A) Overall view of one ATP-cone (green) showing the inner and outer sites binding AMPPNP (proxy for ATP) and dATP respectively. The second ATP-cone in the dimer is shown in beige. The Mg^2+^ ion is drawn as a green sphere. 2m|Fo|-D|Fc| electron density for the nucleotides contoured at 1.0 σ is shown as a grey mesh. B) Details of the interactions between the nucleotides and the ATP-cone. The orientation is approximately the same as in A). C) Closeup of the dATP molecule in the outer site. A 2m|Fo|-D|Fc| omit electron density for dATP contoured at 1.0 σ is shown as a grey mesh. D) Details of the nucleotide interactions in the tetramer of the ADP-dATP form. The interactions are identical to within experimental error in the other chain, which forms infinite “filaments” in the crystal.

As is typical for ATP-cones, the negative charges on the di/triphosphate tails are neutralised by a Mg^2+^ ion bound between them and by a number of Arg and Lys side chains, one of which (Lys76) comes from the second ATP-cone of the pair. Of the side chains, Lys53 plays an important role, as it makes H-bonds to the β-phosphate group of ATP, the γ-phosphate of dATP and also to N7 on the base of ATP. As no other NrdR side chain makes contact with both nucleotides, this is consistent with an observation that the K53A mutation eliminates binding of a second nucleotide to the site when already occupied by either ATP or dATP (13). The authors concluded that ATP and dATP bind with negative cooperativity; however, the structures show that the cooperative effect involves two distinct nucleotide binding sites and not a single site as proposed previously (13). In *S. coelicolor,* single amino acid substitutions K50A (K53A in EcoNrdR), V48A (V51), E56A (E59) and K62A (K65) reduced nucleotide binding, altered NrdR oligomeric state and impaired its ability to bind DNA. The first three of these mutations contact the inner site ATP and the latter contacts the ɣ-phosphate group of the outer site dATP. Interestingly, the Y128A mutant of ScoNrdR (Y131 in EcoNrdR) bound two dATP molecules per ATP-cone instead of one ATP and one dATP (12).

To investigate the thermodynamics of binding of ATP, ADP and dATP to NrdR, we used isothermal titration calorimetry (ITC). All three nucleotides bind with similar affinities to their respective sites in NrdR (K_D_ = 343, 346 and 610 nM for ATP, ADP and dATP, respectively) and the binding curves are consistent with a single set of binding sites for each ligand (Fig. 7). The interactions are enthalpy-driven, with ΔH values of −60, –51 and −57 kJ/mol for ATP, ADP and dATP, respectively (Fig. 7B). The fitted apparent N value slightly below 0.5 could mean that only 50% of the protein in the sample is active due to its relative instability. The crystal structures, which used higher nucleotide concentrations, support a 1:1 binding stoichiometry, and our ITC results suggest that each nucleotide has a strong affinity to only one of the two potential binding sites.

**Figure 7.**
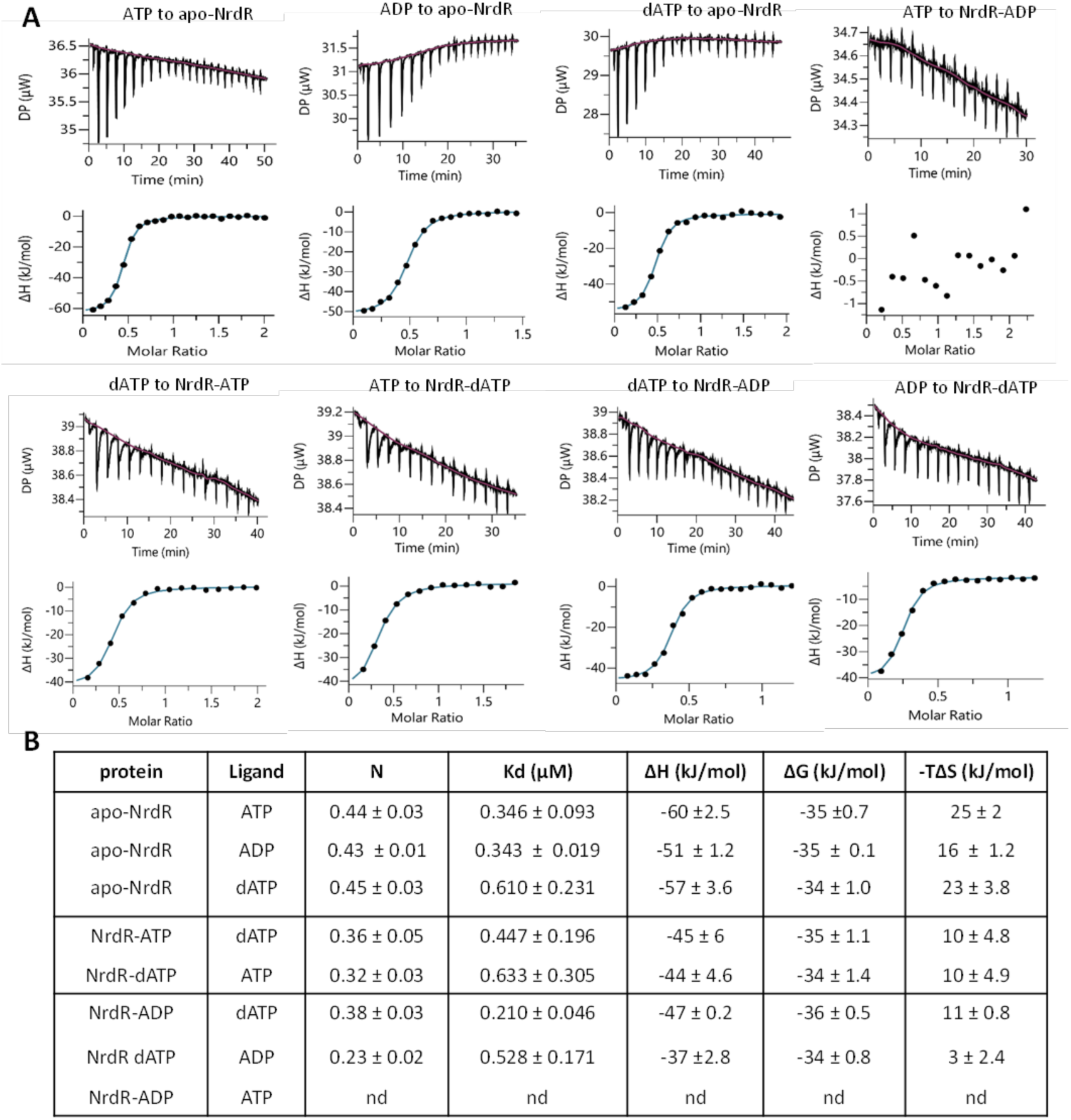
ITC analyses of ligand binding to *E. coli* NrdR at 10°C. (**A**) Representative ITC thermograms obtained by titration of ATP, ADP or dATP to apo-NrdR and titration of ATP, ADP or dATP to NrdR loaded with ATP, ADP or dATP (see Materials and Methods for details). Isothermal calorimetric enthalpy changes (upper panels) and resulting binding isotherms (lower panels) are shown. **(B)** Thermodynamic parameters of ligand binding to NrdR. Binding isotherms were fitted using a one-set-of-sites binding model. Values are reported as the mean ±SD of three titrations. The titrations were performed as described in Materials and Methods. nd, not detected.

To investigate binding of two different effectors, we studied binding of one nucleotide to NrdR preloaded with another nucleotide. The nucleotide concentration in the syringe was adjusted to be insufficient to displace the preloaded nucleotide, if both were competing for the same binding site. Titrating dATP into ATP-loaded NrdR gave a K_D_ of 447 nM and titrating ATP into dATP-loaded NrdR gave a K_D_ of 633 nM (Fig. 7). Titrating dATP to an ADP-loaded sample and vice-versa gave K_D_ values of 210 nM and 525 nM, respectively. The enthalpies of interaction (ΔH values) in this experimental setup decreased by 10-15 kJ/mol, and the N values were slightly lower (Fig. 7B). No binding was detected when ATP was titrated into ADP-loaded NrdR (Fig. 7). These results show that ATP and ADP can bind to the same site whereas dATP binds to another site on NrdR, which is confirmed by the crystal structures. The K_D_ for dATP was essentially the same whether NrdR was unloaded or filled with ATP or ADP, and the same K_D_s were observed for ATP and ADP respectively to both dATP-loaded and unloaded NrdR (Fig. 7B). Our ITC results show that ribonucleotides and deoxyribonucleotides bind independently to different sites.

Since apo-NrdR and ATP-loaded NrdR have decreased stability at higher temperatures, titrations were performed at 10 °C, to avoid the risk of aggregation or precipitation of the sample during the course of the experiment. To confirm the results obtained at 10°C we also performed titration of ATP into apo-NrdR and titration of dATP into ATP-loaded NrdR at 20 °C, 25 °C and 30°C (Supplementary Fig. S15). Our results agree with those of McKethan and Spiro (13), who investigated binding of nucleotides to NrdR, but incorrectly interpreted their results as binding of nucleotides to a single binding site with negative cooperativity.

### Cryo-EM structure of ATP-loaded EcoNrdR filaments

In the presence of ATP alone, EcoNrdR formed extended filaments on the cryo-EM grids. The structure of a portion of the filaments was determined to approximately 6.4 Å resolution. ATP-cone and Zn-ribbon pairs from the AMPPNP-dATP complex were fitted as rigid bodies into the map, with excellent agreement (Fig. 8A). The helical filaments consist of tetramers similar to those in the AMPPNP-dATP complex, but flatter and more elongated (Fig. 8B). The relative orientations of the ATP-cone and Zn-ribbon domains in the ATP-only filaments are very similar to those in the ATP-dATP tetramers, having two distinct conformations that fall into the clusters described earlier (Fig. 4C and Supplementary Figure S11). In contrast, there is a significant rotation of the ATP-cone domains relative to each other (Supplementary Movie S4). This is similar to the significant shifts in ATP-cone orientation observed between the ATP- and ATP-dATP-bound forms of ScoNrdR. In that system, the nucleotide bases stack on top of each other in the ATP-bound dodecamer and are ∼8 Å closer to each other than they are in the ATP-dATP form. In contrast, in EcoNrdR they do not approach each other as closely. However, this analysis assumes that the nucleotides move as a rigid body with the rest of the ATP-cone. A more detailed analysis of differences between the ATP-bound filament of tetramers and the AMPPNP-dATP-bound tetramers requires a higher resolution cryo-EM reconstruction.

**Figure 8.**
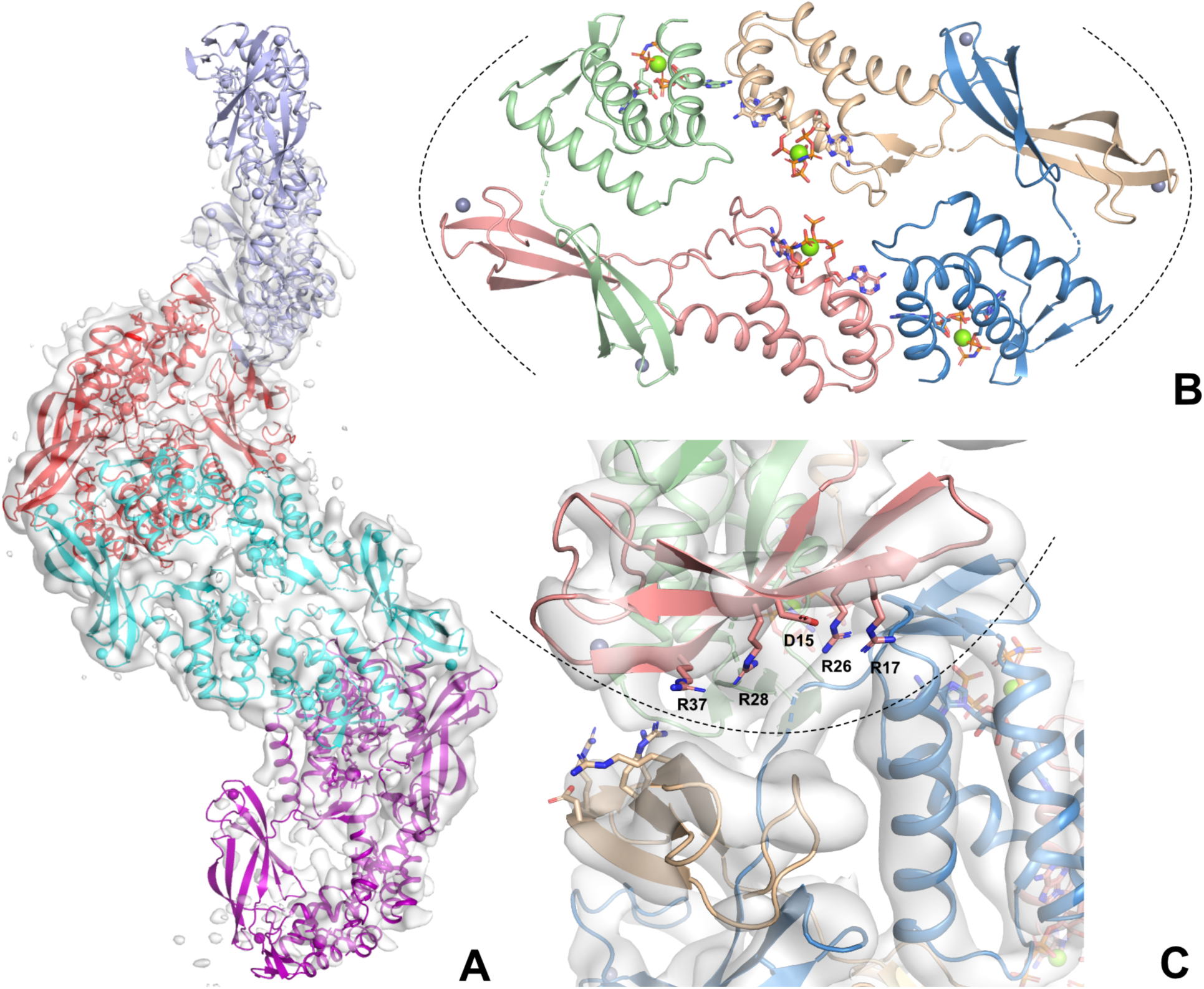
Cryo-EM structure of the EcoNrdR filaments formed in the presence of ATP. A) Four tetramers from the filament. The cryo-EM map with a nominal resolution of 6.4 Å is shown contoured at 7.0 σ above the mean. At this contour level the central two tetramers (red and cyan) are very well defined but the outer two (lilac and purple) are only visible at lower contour levels. B) Structure of one of the central tetramers (red in panel A). The four chains of the tetramer are coloured as in previous figures. Nucleotides are shown as sticks. C) Closeup view of the interface between the two middle tetramers in panel A. Residues involved in DNA recognition are shown as sticks. The dotted line indicates the interface regions shown in panel B.

The interface between tetramers in the helix involves the ATP-cones of one monomer and the Zn-ribbon of another (see the dotted lines in Fig. 8B). While the resolution of the cryo-EM reconstruction precludes a detailed analysis of the interactions between side chains, rigid body fitting of the domains to the map suggests a plausible hydrophobic core with several additional salt bridges between tetramers. Interestingly, the face of the Zn-ribbon pair that hosts the conserved residues involved in DNA recognition (Asp15, Arg17, Arg26, Arg28 and Arg47) is sequestered in the interface between tetramers and thus unavailable for DNA binding. This is similar to the burial of the DNA binding residues in the ATP-bound dodecamer of ScoNrdR (1).

## Discussion

The transcriptional regulator NrdR was first discovered in *S. coelicolor* (2) and later extensively studied in *E. coli* and *P. aeruginosa* (10,13,14,16,33). While these studies were performed with methods available at the time, we have now used a combination of highly specific and sensitive methods to study the prerequisites of the regulatory mechanism of NrdR in *E. coli*, including the first structural studies of EcoNrdR. We have identified a new NrdR binding site in the *nrdAB* promoter region comprising one previously known NrdR box (10) and one new NrdR box (this study), and subsequently demonstrated that all three *E. coli* RNR operons (*nrdAB, nrdHIEF, nrdDG*) bind NrdR equally well. Notably, additional transcription factors bind RNR promoters: DnaA and H-NS to *NrdA*, Fur and IscR to *NrdHEF*, H-NS and Fnr to *NrdDG* (7), making the regulation a fine-tuned interplay between NrdR, other regulators and RNA polymerase.

In general, the DNA-binding forms of NrdR loaded with dATP plus ATP or ADP are tetrameric in solution, whereas the non-binding form loaded with ATP forms higher oligomers, and apo-NrdR is dimeric. We have determined crystal structures of the EcoNrdR tetramer in complex with dATP and ATP (or its analogue AMPPNP), as well as in complex with dATP and ADP. Furthermore, we have determined cryo-EM structures of the ATP-dATP tetramer bound to DNA and of an inactive filament of EcoNrdR formed in the presence of ATP only.

A key observation is that the EcoNrdR tetramer displays considerable flexibility even in the presence of a specific nucleotide combination. In the AMPPNP-dATP and ATP-dATP forms, the orientations of the Zn-ribbon domains relative to the ATP-cones fall into two distinct clusters in the 12 individual monomers, with considerable variation within the clusters. This intrinsic flexibility seems to be a prerequisite for DNA binding. Supplementary Movie S5 shows hypothetical models of an encounter of one of the Zn-ribbon pairs with one of the NrdR boxes in undistorted B-DNA, followed by remodelling of both DNA and protein to allow interaction of the second Zn-ribbon pair with its NrdR box. Such a transition is also possible for the ADP-dATP form, which is shown in the biophysical assays to have the same affinity for DNA as the ATP-dATP form. However, we cannot exclude that the ADP-dATP form in solution is more similar to the ATP-dATP form than the crystal structure.

There are several striking differences between the structures of EcoNrdR and the previously determined NrdR from *S. coelicolor* (ScoNrdR) (1). Firstly, the dATP-ATP bound ScoNrdR is an octamer that can dissociate to and is in equilibrium with a tetramer, whereas EcoNrdR under comparable conditions is a tetramer that oligomerizes at increasing protein concentrations.

Secondly, the conformation of the dATP-ATP loaded ScoNrdR tetramer is very similar in the DNA-bound and unbound states, whereas the relative orientations of the ATP-cones and Zn-ribbons in the dATP-ATP loaded EcoNrdR tetramer change significantly upon DNA binding (Figure 4C, 4D and Supplementary Movie S2). This may relate to the fact that EcoNrdR seems more prone to accept variations in distance between the NrdR boxes *in vitro* compared to ScoNrdR (34). The linker between the two protein domains in EcoNrdR is three residues longer, which may allow greater flexibility. Thirdly, ATP-loaded EcoNrdR forms filaments both in solution and on cryo-EM grids, whereas the ATP-loaded ScoNrdR was a compact dodecamer under all conditions tested. As the DNA-binding faces of the Zn-ribbons are buried in tetramer-tetramer interactions within the filament (Figure 8C), the filaments likely represent an alternative mechanism for sequestration of NrdR in an inactive form. The conformation of EcoNrdR in the monomers of the filament does not strongly resemble that in the ScoNrdR dodecamers, suggesting that ATP acts by a different molecular mechanism in the two organisms. Interestingly, SAXS data of *Acinetobacter baumannii* NrdR suggests formation of filaments, which dissociate in the presence of DNA adenine methyltransferase (35).

With the high resolution of the EcoNrdR crystal structures we can describe the two nucleotide binding sites in the ATP-cone in detail. Our structural and ITC results confirm that binding to the ‘inner’ and ‘outer’ sites occurs independently. Interestingly, loss of the γ-phosphate group in the ADP-dATP complex results in only minimal perturbation of the ATP cone structure, in particular the relative orientations of the ATP-cone pairs are identical (Figure 6B,D). In contrast, the ATP-cone and Zn-ribbon domains have a very different relative orientation to those in the AMPPNP-dATP or ATP-dATP complexes. The ADP-dATP form may be to some extent a result of crystal packing effects. In particular, it is unclear why the crystal consists of tightly packed tetramers combined with infinite linear chains of monomers linked through their ATP-cone and Zn-ribbon domains. Nevertheless, all these structures are compatible with a model in which EcoNrdR complexes in the ATP(ADP)-dATP form are highly conformationally adaptable (and thus somewhat prone to crystal packing effects).

Our combined studies of EcoNrdR and ScoNrdR clearly demonstrate a physiological mechanism depending on the simultaneous binding of dATP plus ATP (or ADP) for repression of RNR operons. An earlier study suggested that EcoNrdR’s DNA binding capacity was more sensitive to phosphate number than to the absence of a 2’-OH group in dATP, and that dephosphorylation of a bound triphosphate to monophosphate promoted DNA binding (13).

However, our MST results clearly show that adenosine monophosphate nucleosides do not trigger DNA binding alone or in combination with another nucleotide. The crystal structures also explain why AMP fails to promote DNA binding while ADP does. The two nucleotides ATP and dATP are tightly intertwined through their triphosphate tails, assisted by a Mg^2+^ coordinated between them. Lack of a γ-phosphate group in ADP results in the loss of only one Mg^2+^ coordination and one H-bond to Lys76. Removal of the β-phosphate group is predicted to result in more far-reaching consequences, as it also coordinates Mg^2+^ but also makes three H-bonds, including one to Lys53, which bridges to the adenine base of ATP and the γ-phosphate of dATP. Thus, the β-phosphate appears critical for binding of Mg^2+^ and for proper orientation of the nucleotides. The critical interactions are maintained in the ADP-dATP complex. Interestingly, EcoNrdR still forms tetramers in the presence of AMP+dAMP, while the apo-form exists as dimers. This suggests that the β- and γ-phosphate interactions are not strictly necessary for association of the ATP-cone domains, though they are critical to achieve the functional repressor state.

The four different conformations of the EcoNrdR tetramer presented in this work suggest a structural basis for modulation of repressor activity by ATP and dATP. In the presence of dATP and ATP (or ADP) there is considerable potential for flexibility between the ATP-cone and Zn-ribbon domains, which in themselves behave as rigid bodies, moving in pairs. The DNA complex reveals a tetramer like the one seen previously for ScoNrdR, in which each pair of Zn ribbons is bound to a NrdR box, but with a less compact form than for ScoNrdR. In contrast to ScoNrdR, the conformational adjustments between free and DNA-bound tetramers are larger in EcoNrdR. In all tetramers where dATP is bound, the ATP-cones have the same relative orientation. In contrast, during transition towards the tetramers of the ATP-loaded filament, the ATP-cones move relative to each other. The ATP-cones and Zn-ribbons have yet another relative orientation, unique to the ATP-bound form. It is possible that only in this form are the two domains and the linker between them (which is part of the interface between tetramers) in a conformation that permits formation of a binding pocket that sequestrates the DNA-binding faces of the DNA-ribbon domains.

A burial of the DNA-binding residues was also seen in the ATP-bound dodecamer of ScoNrdR, which consisted of three tightly interleaved tetramers. However, the movement of ATP-cone domains between ATP-dATP and ATP-bound forms in ScoNrdR is different. It is clear that the ATP molecules are further from each other in ATP-bound EcoNrdR, while they approach each other and the bases stack on each other in the equivalent form of ScoNrdR. The low resolution of the ATP-bound EcoNrdR filament precludes a more detailed analysis of how the very small difference which is the addition of a 2’-OH group in ATP can cause the sequestration of EcoNrdR in a filamentous form, and it will be an important goal of future studies to analyse this at higher resolution.

## Materials and Methods

### Plasmids

The *nrdR* gene from *E. coli* K-12 strain MG1655 was amplified and cloned into the pET30a(+) vector (Novagen) as described previously (10) resulting in pET30a(+)::*nrdR* construct expressing wild type NrdR protein (NP_414947.1) with a C-terminal hexahistidine (His) tag.

### Reagents

ATP and dATP were purchased from ThermoFisher Scientific as 100 mM solutions at pH 7.0. Other nucleotides were purchased from Sigma Aldrich, stock solutions of 50-250 mM were prepared in water and the pH was adjusted to 7. AMPPNP and c-di-AMP were obtained from Jena Bioscience, Germany in lyophilized form and resuspended in water to obtain a 10 mM solution.

### Protein expression

Overnight cultures of *E. coli* BL21(DE3) bearing pET30a(+)::*nrdR* were diluted to an absorbance at 600 nm (OD_600_) of 0.1 in LB (Luria-Bertani) liquid medium, containing kanamycin (50 μg/ml) and shaken vigorously at 37 °C. At OD_600_ = 1, 0.4 mM isopropyl-β-D-thiogalactopyranoside (Sigma) and 0.1 mM Zn(CH_3_CO_2_)_2_ were added. The cells were grown overnight under vigorous shaking at 20 °C and harvested by centrifugation. The cell pellet was stored at –80 °C.

Selenomethionine-derivatized NrdR was produced by transforming the pET30a(+)::*nrdR* plasmid into the auxotrophic strain *E. coli* B834 (DE3). Cells were cultured in SelenoMet Minimal Media (Molecular Dimensions Ltd), containing kanamycin (50 μg/ml), according to the manufacturer’s instructions, with the exception that seleno-L-methionine (Sigma Aldrich) was used at 40 mg/l.

### Protein purification

The cell pellet was thawed and resuspended in lysis buffer: 50 mM Tris-HCl pH 8.5 (at 4 °C) containing 300 mM NaCl, 10 mM imidazole and 2 mM DTT or 0.2 mM Tris(2-carboxyethyl)phosphine (TCEP). Phenylmethylsulfonyl fluoride (PMSF) was added at 1 mM to the cell suspension, the mixture was sonicated in an ultrasonic processor (Misonics) until clear and the lysate was centrifuged at 18,000 × g for 45 min at 4 °C. The supernatant was loaded on a HisTrap FF Ni-Sepharose column (Cytiva) on an ÄKTA prime system (Cytiva), equilibrated with lysis buffer (without PMSF), washed thoroughly with buffers containing 10 mM and 60 mM imidazole and eluted with buffer containing 500 mM imidazole. The proteins were desalted using a HiPrep 26/10 Desalting column (Cytiva) in 50 mM Tris-HCl pH 8.6 at 4°C, 300 mM NaCl and 1 mM TCEP, frozen and stored at −80 °C. The zinc content of NrdR was 0.67 ± 0.01 mol Zn/mol protein (based on three independently prepared samples) quantified using total-reflection X-ray fluorescence (TXRF) on a Bruker PicoFox S2 instrument and analysed with software provided with the spectrometer. A gallium internal standard at 2 mg/l was added to the samples (v/v 1:1) before the measurements.

To remove bound nucleotides the proteins were applied to a HiTrap Phenyl FF (high sub) column (Cytiva) in 50 mM Tris-HCl, pH 8.6 and 0.7 M (NH_4_)_2_SO_4_ and 0.1 mM TCEP, washed extensively (70 column volumes) with the same buffer, washed with buffer without ammonium sulphate (most of the protein remained bound to the column at this stage) and eluted with water with 0.1 mM TCEP. Protein recovery after this stage was approximately 20%. Buffer components were added to the sample eluted with water. Buffer normally contained 50 mM Tris-HCl pH 9.0 (at 4°C), 300 mM NaCl and 1 mM TCEP. Buffer for the NrdR sample used for microscale thermophoresis (MST) was at pH 8.6 and also contained 20 mM MgCl_2_. For the NrdR sample used for ITC, buffer exchange was performed using HiPrep 26/10 Desalting column equilibrated with 50 mM Tris-HCl pH 9.0 (at 4°C), 300 mM NaCl and 1-2 mM TCEP. Protein concentration was determined using Coomassie Plus (Bradford) Assay Kit (ThermoFisher Scientific) using a BSA standard curve. Zinc content of apo-NrdR was 0.63 ± 0.05 mol Zn/mol protein (based on three independently prepared samples) quantified using TXRF as described above. Nucleotide content of apo-NrdR was ∼0.2 mol nucleotide/mol protein quantified after denaturation of NrdR in 1 M HCl, centrifugation and quantification of the nucleotide in the supernatant at A_260_ using extinction coefficient 15.4 x 10^3^ AuM^-1^. The apo-NrdR was aliquoted, frozen in liquid nitrogen and stored at −80°C until used at concentrations of approximately 0.9 - 1.8 mg/ml. The apo-NrdR cannot be further concentrated without addition of effector nucleotides due to apo-protein instability at higher concentrations.

For crystallisation of NrdR with ATP and dATP, MgCl_2_, ATP and dATP were added to apo-NrdR to reach 6, 0.6 and 0.6 mM respectively. The sample was incubated for 20 minutes at 8 °C and applied to a Superdex 200 pg 16/600 column equilibrated with buffer containing 25 mM Tris-HCl pH 9.0 (at 4 °C), 300 mM NaCl, 1 mM TCEP, 10 mM MgCl_2_, 0.1 mM ATP and 0.1 mM dATP.

Fractions from the middle of the single eluted peak were collected. ATP and dATP were added to 1 mM each and glycerol was added to 5%. The sample was then concentrated to 13.5 mg/ml using Vivaspin 500 concentrators with 50,000 molecular weight cutoff (MWCO) PES membranes (Sartorius) on a tabletop centrifuge at 4 °C.

For crystallisation of NrdR in the presence of ADP and dATP or with AMP-PNP and dATP, NrdR samples after nickel affinity chromatography and desalting were diluted in 1:1 ratio in SEC buffer (see below) and supplemented with either 1.5 mM ADP and 1.1 mM dATP or 1.5 mM AMP-PNP and 1 mM dATP, incubated at 8 °C for 15 minutes and applied to a Superdex 200 Increase 10/300 GL column equilibrated with SEC buffer containing 25 mM Tris-HCl pH 9 (at 4°C), 300 mM NaCl, 1 mM TCEP and 10 mM MgCl_2_. SEC buffers used for ADP and dATP-NrdR contained 0.2 mM ADP and 0.2 mM dATP. No nucleotides were included in the SEC buffer used for the sample with AMP-PNP and dATP, which instead was supplemented with 0.5 mM AMP-PNP and 0.5 mM dATP after elution.

For crystallisation of NrdR grown in the presence of seleno-methionine the protein was supplemented with 0.1 mM zinc acetate 10 mM MgCl_2_, 3 mM ATP and 1 mM dATP, incubated for 15 minutes at 8 °C, concentrated using Amicon Ultra-15 centrifugal filters (Ultracel-50K, Merck Millipore) and applied to a Superdex 200 Increase 10/300 GL column equilibrated with SEC buffer containing 25 mM Tris-HCl pH 8.5, 300 mM NaCl, 1 mM TCEP, 10 mM MgCl_2_, 0.2 mM ATP and 0.2 mM dATP.

All protein samples eluted as one peak and the fractions from the middle of the peak were collected and concentrated using Vivaspin 500 (50 000 MWCO PES) on a tabletop centrifuge at 4 °C to 17 mg/ml and 8 mg/ml for samples containing ADP and dATP and AMP-PNP and dATP respectively and to 10 mg/ml for selenomethionine ATP- and dATP-bound NrdR.

### Protein crystallisation and data collection

All crystals were produced using sitting drop vapour diffusion using a mosquito robot (SPT Labtech). Wild type apo-EcNrdR loaded with ADP and dATP at 13.5-17 mg/mL in buffer as described above was crystallised by mixing 150 nL protein with 50 nL reservoir solution containing 0.1 M Bis-Tris pH 5.5, 0.2 M ammonium acetate, 45% (v/v) methane pentane diol at 20 °C. The crystals obtained were directly harvested and flash-frozen in liquid nitrogen. X-ray data were collected at 100 K and wavelength 0.9795 Å on the I04 beamline at Diamond Light Source, Didcot, UK.

Wild type Apo-EcNrdR loaded with AMPPNP and dATP at 8 mg/mL was crystallised by mixing 150 nL protein with 50 nL reservoir solution containing 0.1 M Bis-Tris propane pH 7.5, 0.2 M potassium thiocyanate, 20% (w/v) polyethylene glycol 33550 (PEG3350) at 20 °C. The crystals obtained were harvested in paraffin oil and flash-frozen in liquid nitrogen. X-ray data were collected at 100 K and 0.9763 Å wavelength at beamline BioMAX of the MAX IV synchrotron, Lund, Sweden.

Selenomethionine-containing Apo-EcNrdR (SeMet-EcNrdR) at 10 mg/mL was crystallised by mixing 100 nL protein with 100 nL reservoir solution containing 0.1 M HEPES pH 7.5, 0.2 M MgCl_2_, 12.5% (w/v) PEG3350 at 20 °C. The crystals obtained were cryoprotected by soaking in mother liquor containing the reservoir supplemented with 30% ethylene glycol. Diffraction data were collected at 100K at BioMAX (wavelength 0.9791 Å).

Data were indexed, processed, and scaled using the autoPROC pipeline and any anisotropy in the diffraction was corrected by STARANISO (17). *S. coelicolor* NrdR (PDB entry 7P3F), was used as a model to solve the structure of the ADP-dATP form by domain-wise molecular replacement using the program Phaser (18). Buster (19) or Phenix (20) were used for refinement and Coot for model reconstruction. The individual domains of the ADP-dATP form were then used to solve the other structures in a similar manner.

### Cryo-EM of the EcNrdR-ATP-dATP-DNA complex

To obtain NrdR bound to DNA, oligonucleotides containing the NrdR-boxes in the promoter region of *nrdEF* with 7 and 3 nucleotides overhang (in bold) on the termini of the sense and antisense oligonucleotides, respectively: EcoEF7,3_For: 5’-**GCACTAT**TTGCTATATATTGTGTGGTTGAATCTTTTTTCAACTACATCTAGTAT**CTC**-3’ and EcoEF7,3_Rev: 5’-**GAG**ATACTAGATGTAGTTGAAAAAAGATTCAACCACACAATATATAGCAA**ATAGTGC**-3’ were ordered from Sigma-Aldrich, resuspended and annealed in buffer containing 10 mM Tris pH 7, 100 mM NaCl and 10 mM MgCl_2_ as described in the Microscale thermophoresis section to obtain EcoEF7,3 double stranded oligonucleotide.

Apo-NrdR was supplemented with nucleotides (either ATP or ADP), dATP and MgCl_2_, mixed with EcoEF7,3 ds oligonucleotide resulting in a sample used for cryo-electron microscopy (cryo-EM) containing 0.31 mg/ml NrdR (16 μM monomeric protein) and 6 μM oligonucleotide in 25 mM Tris-HCl pH 8.5 (at 4 °C), 100 mM NaCl, 5 mM MgCl_2_, 1 mM TCEP, 0.5 mM dATP and 0.5 mM ATP/ADP.

For cryo-EM grid preparation, 4 μl sample was applied to each glow-discharged (30 s at 15 mA) QuantiFoil 2/1 Cu 300 grid (Quantifoil Micro Tools GmbH), pre-coated with graphene oxide according to Boland et al (21). Grids were vitrified in liquid ethane using a Vitrobot Mark IV (Thermo Fisher Scientific) at 100% humidity, 4 °C, using a blot force of −5, wait time of 1 s, and blotting time of 5 s. Automated data collection was performed at the Umeå Centre for Electron Microscopy using EPU software on a Titan Krios G2 transmission electron microscope (Thermo Fisher Scientific) operated at 300 kV, equipped with a Gatan K2 direct electron detector and BioQuantum energy filter. Two datasets were collected at 0° and 30° stage tilt respectively, using a pixel size of 0.63 Å, total dose of 58 e^−^/Å^2^, and a defocus range of −1.2 to −2.4 μm.

After motion correction, contrast transfer function (CTF) correction and curation, the untilted and tilted datasets contained 7472 and 7317 micrographs respectively. The final reconstruction used both datasets. The procedure is summarised in Supplementary Figure S1. Briefly, particles were picked from 500 micrographs from each dataset using the blob picker in CryoSPARC v4.2.1 (22). The 188 697 particles extracted thereafter were twice classified into 50 classes, of which nine (24 351 particles) were selected for a template that was used to pick particles from a set of 10 787 micrographs. Around 3.4 M extracted particles (box size 512 pixels) were classified into 100 2D classes, of which 15 (964 338 particles) were used to make three ab-initio models. These were all subjected to homogeneous and non-uniform (NU) refinement. The best model had a resolution of 4.65 Å. The 379 199 particles from this model were once again 2D-classified to remove junk particles, reducing the number to 178 716. A new *ab initio* model was made from these, followed by homogeneous and NU-refinement with C2 symmetry, giving a final resolution of 4.06 Å. As the DNA-bound EcoNrdR tetramer is relatively small, the initial lowpass filtering resolution was adjusted to 15 Å. The dynamic mask parameters were altered to near: 1.5 Å and far: 14 Å, resulting in tighter dynamic masking. The final reconstruction was locally sharpened using DeepEMhancer (23).

Three models were used to fit the best cryo-EM volume: the *S. coelicolor* DNA from the ScNrdR-dATP-ATP-DNA complex, and pairs of ATP-cones and Zn-ribbons from the EcNrdR-ATP-dAMPPNP complex. These were fitted to the volume using Molrep (24) from the CCP-EM package (25) in the order DNA:ATP-cones:Zn ribbons. Rigid body real space refinement was then carried out using phenix.refine (26). In the final step, the ATP-cones in each pair were refined independently, but they did not change orientation significantly relative to each other.

### Cryo-EM of EcNrdR-ATP filaments

For the ATP-bound sample, apo-NrdR was supplemented with 300 μM MgCl_2_ and 300 μM ATP and applied to a SEC column equilibrated with buffer containing 50 mM Tris-HCl, pH 9 (at 4 °C), 300 mM NaCl, 5 mM DTT, 0.2 mM MgCl_2_ and 0.1 mM ATP. Eluted NrdR at concentration of 0.2 mg/ml (corresponding to 10 μM monomeric NrdR) was applied to Quantifoil R2/2 Cu 300 mesh grids, previously coated with graphene oxide using a 0.2 mg/ml suspension. Sample vitrification was carried out using a Vitrobot Mark IV. 1,672 cryo-EM movies were collected at the Swedish National Cryo-EM Facility in Stockholm using a Titan Krios microscope equipped with a K3 direct electron detector and a Bioquantum energy filter (Gatan). The data acquisition parameters and data processing workflow are presented in Supplementary Figure S2. Data processing including patch motion correction, particle picking, 2D classification and 3D refinement, carried out in CryoSPARC v2.1 (22), CTF estimation was performed using CTFFIND4 (27) and model building was carried out in Coot v0.9.8 (28). Four copies of the EcoNrdR-ATP-dATP tetramer were placed in the cryo-EM volume, which is of excellent quality for the central two tetramers but less well defined for the outer two. Domain-wise rigid body real space refinement was then carried out in Coot. No automatic refinement was done.

### Microscale thermophoresis

We studied the binding of *E. coli* NrdR to its cognate sites in the promoter regions of *nrdAB* (RNR class Ia), *nrdEF* (RNR class Ib) and *nrdD* (RNR class III) comprising two 16 bp palindromic sequences separated by 15-16 bp, termed NrdR-boxes (11), using MST (29). Oligonucleotides (57-58 bases) containing the NrdR boxes (underlined) and flanking regions of 5 nucleotides on each side, and a negative control *E. coli ribX* gene promoter were ordered from Genscript. The sense oligonucleotides were fluorescently labelled by the Cy5 dye at their 5’ end by the manufacturer. For *nrdAB* (nrdAB_0,1_Cy5_sense: 5’-Cy5-CATTT<u>TCACACTATCTTGCAG</u>TGAATCCCAAACATAC<u>CCCCTATATATAGTGT</u>TCTAA-3’, *nrdAB_*0,1_antisense: 5’-TTAGAACACTATATATAGGGGGTATGTTTGGGATTCACTGCAAGATAGTGTGAAAATG-3’ and nrdAB_1,2_sense: 5’-Cy5-CATACCCCCTATATATAGTGTTCTAAGCAGCTTCCCGTACTACAGGTAGTCTGCATG-3’, nrdAB_1,2_antisense: 5’-CATGCAGACTACCTGTAGTACGGGAAGCTGCTTAGAACACTATATATAGGGGGTATG-3’) *nrdEF* (nrdEF_Cy5_sense: 5’-CY5-ACTAT<u>TTGCTATATATTGTGT</u>GGTTGAATCTTTTTT<u>CAACTACATCTAGTAT</u>CTCTG-3’, *nrdEF*_antisense: 5’-CAGAGATACTAGATGTAGTTGAAAAAAGATTCAACCACACAATATATAGCAAATAGT-3’), *nrdDG* (nrdRDG_Cy5_sense: 5’-CY5-GCAAA<u>GCACTATATATAGACT</u>TTAAAATGCGTCCCA<u>ACCCAATATGTTGTAT</u>TAATC-3’, *nrdDG*_antisense: 5’-GATTAATACAACATATTGGGTTGGGACGCATTTTAAAGTCTATATATAGTGCTTTGC-3’) and *ribX* (ribX_Cy5_sense: 5’-CY5-AGCATCGATCCTTTATCTCAAAAGCGTTGCGCCTTTGTTGTATCGTCAGTTCAGGGT-3’, ribX_antisense: 5’-ACCCTGAACTGACGATACAACAAAGGCGCAACGCTTTTGAGATAAAGGATCGATGCT-3’). Freeze dried oligonucleotides were reconstituted in Tris 50 mM pH 8, NaCl 50 mM and EDTA 1 mM. 50 and 57.5 pmol of labelled and unlabeled oligonucleotides respectively were mixed in a total volume of 50 µl containing 50 mM Tris-HCl pH 7.4, 50 mM NaCl and annealed using a thermoblock. The annealing program included incubation for 5 min at 95 °C and gradual cooling to 25 °C using 140 cycles of −0.5 °C and 45 s per cycle, resulting in 1 µM double-stranded DNA.

MST was performed using a Monolith NT.115 instrument (NanoTemper Technologies GmbH). Binding was measured between either apo- or as-purified NrdR and double stranded oligonucleotides containing the promoter region of *nrdAB, nrdEF, nrdDG* and *ribX* labelled with Cy5 in MST buffer containing 17 mM Tris-HCl pH 8.6, 100 mM NaCl, 7 mM MgCl_2_, 0.33 mM TCEP, 0.025% Tween-20 and the specified nucleotide or a combination of nucleotides (1 mM each, unless otherwise stated) at room temperature. We verified that the residual nucleotides in as-purified NrdR were replaced by either ATP and dATP or ADP and dATP after their addition to the sample (see Supplementary Figure S3 and its legend). A 16-step dilution series was prepared by adding 10 μL buffer to 15 tubes. NrdR, 20 μL of 40 - 90 μM protein (at least three different protein preparations were used in MST experiments), was placed in the first tube, and 10 µL was transferred to the second tube and mixed well by pipetting (1:1 dilution series). To each tube of the dilution series 10 µL of the binding partner (double stranded Cy5 labelled oligonucleotide in the same buffer) was added to reach 5 nM. The samples were incubated for 10 minutes at room temperature before being transferred to Monolith™ NT.115 Series Capillaries (NanoTemper). The MST measurements were performed using high MST power and 50% laser excitation power. The results were analysed using the MO.Affinity Analysis v2.3 software (Nanotemper) with default parameters. For dATP + ATP and dATP + ADP nucleotide combinations, K_D_ and standard deviation were calculated using fits from at least three individual titrations. In individual cases, such as the combination of nucleotides AMP and dATP (binding to *nrdAB, nrdEF* and *nrdDG* promoters), as well as in the presence of ADP (*nrdEF* promoter), dATP (*nrdEF* and *nrdDG* promoter) and the combination of c-di-AMP and ADP (*nrdDG*), the K_D_s could not be reliably determined since the titration curves did not reach a plateau (see titration curves in Supplementary Materials). The indicated K_D_ values in the high micromolar range derived from these curves (Table 1) are therefore only estimates.

### Analytical size exclusion chromatography

To study nucleotide dependent protein oligomerization, SEC experiments were performed at a temperature of 4-8 °C using a Superdex 200 PC 3.2/30 column (with a total volume of 2.4 ml) connected to a Shimadzu high pressure liquid chromatography (HPLC) system, alternatively to the ÄKTA prime system (Cytiva). The column was equilibrated with degassed SEC buffer containing 25 or 50 mM Tris-HCl pH 9.0 at 4 °C, 300 mM NaCl, 0.2-10 mM MgCl_2_, 2-5 mM dithiothreitol (DTT) or 0.5 mM TCEP and when indicated, ATP, ADP or combinations of ATP and dATP, ADP and dATP or AMP and dAMP at 0.1-0.5 mM each. Nucleotides were included in the SEC buffer to avoid dissociation of nucleotide-induced protein complexes during the run. In experiments in which the ATP-loaded NrdR sample eluted in the resolution range of the column, the concentration of MgCl_2_ in the SEC buffer was 0.1-0.2 mM. When MgCl_2_ in SEC buffer was 1 mM or higher, NrdR eluted at the void volume of the column. For NrdR samples loaded with other effectors, the concentration of MgCl_2_ did not affect the oligomeric state of NrdR. The flow rate was 0.1 ml/min on Shimadzu HPLC system and 0.7 ml/min on an ÄKTA prime system.

Protein samples for SEC were prepared as follows: combinations of nucleotides ATP and dATP, ADP and dATP, AMP and dAMP premixed with equimolar concentrations of MgCl_2_ were added to apo-NrdR to 0.1-1 mM of each nucleotide and 0.2-10 mM MgCl_2_.

In the sample with ADP the concentrations of ADP and MgCl_2_ were 0.2 mM. In samples with ATP the concentration of ATP was 0.1 mM and MgCl_2_ 0.1 or 0.2 mM. The samples were pre-incubated for 10-60 min at 8 °C, centrifuged and applied to the column. 10-93 μl of 55 μM apo-NrdR, 50 μl of 17-55 μM ATP-loaded NrdR, 25-95 μl of 7-55 μM NrdR loaded with ATP and dATP, 25-93 μl of 10-90 μM NrdR loaded with ADP and dATP were assayed. Molecular weights were estimated based on a calibration curve derived from globular protein standards using both high and low molecular weight SEC marker kits (Cytiva) for each respective column and system. Standard deviations were calculated based on at least three individual chromatograms.

### Isothermal titration calorimetry

ITC experiments were carried out on a MicroCal ITC200 system (Malvern Panalytical) 50 mM Tris-HCl, pH 9, 300 mM NaCl, 1 mM TCEP and 1 mM MgCl_2_. Measurements were done at 10°C as well as at 20 °C, 25 °C and 30 °C, with stirring speed 1000 rpm. The initial injection volume was 0.6-1 μl over a duration of 1.2-2 s. All subsequent injection volumes were 2-2.5 μl over 4-5 s with a spacing of 150-180 s between the injections. Data for the initial injection were not considered. The concentration of NrdR in the cell was 5-20 μM. In the syringe were 50-400 μM ATP, ADP and 75-200 μM dATP. In experiments where NrdR was first preloaded with a nucleotide, an initial ITC titration with one nucleotide was performed, followed by another titration of the second nucleotide into the same sample. Alternatively, the first nucleotide was added to apo-NrdR at a concentration sufficient to saturate the binding site, (based on the results from single nucleotide titrations, normally 1-2 fold the protein concentration) followed by titration with the second nucleotide. Both approaches gave similar results. The data were analysed using the one set of sites model of the MicroCal PEAQ-ITC analysis software (Malvern Panalytical). Standard deviations in thermodynamic parameters, N and K_d_ were estimated from the fits of three different titrations.

## Data availability

The crystal structures presented here are openly available in the Protein Data Bank (www.pdb.org) at https://doi.org/10.2210/pdb9FXK/pdb (AMPPNP-dATP), https://doi.org/10.2210/pdb9FVR/pdb (SeMet-ATP-dATP) and https://doi.org/10.2210/pdb9FZF/pdb (ADP-dATP). The cryo-EM maps for the ATP-dATP-DNA complex and for the DNA filaments are openly available from the Electron Microscopy Data Bank (EMDB; https://www.ebi.ac.uk/pdbe/emdb) with accession numbers EMD-50819 and EMD-50849 respectively. The coordinates fitted to these maps are available on request from the corresponding author.

## Funding

This work was supported by Carl Trygger Foundation [grant number CTS 20:361 to I.R.G.], the Swedish Research Council [grant numbers 2019-01400 to B.-M.S., 2016-04855 and 2023-05074 to D.T.L. and 2022-03681 to P.S], the Swedish Cancer Foundation [grant number 20 1210 PjF to B.-M.S. and 20 1287 PjF to P.S], and the Wenner-Gren Foundation [to B.-M.S.].

## Acknowledgements

Cryo-EM data were collected at the Umeå Core Facility for Electron Microscopy and the SciLifeLab Stockholm cryo-EM facility, which are both nodes of the Cryo-EM Swedish National Facility, funded by the Knut and Alice Wallenberg, Family Erling Persson and Kempe Foundations, SciLifeLab, Stockholm University and Umeå University. We thank Michael Hall for help with data collection. We acknowledge MAX IV Laboratory for time on beamline BioMAX under proposal 20200277 and thank beamline staff for assistance. Research conducted at MAX IV, a Swedish national user facility, is supported by the Swedish Research Council under contract 2018-07152, the Swedish Governmental Agency for Innovation Systems under contract 2018-04969, and Formas under contract 2019-02496. We thank Diamond Light Source for beamtime (proposal mx21625) and the staff of beamline I04 for assistance. We thank Henrietta Nielsen, Stockholm University, for letting us use the MST instrument.

## Conflict of interest

The authors declare no conflict of interest.

## Author contributions

IRG, OB, MMC, PS, BMS and DTL planned experiments. IRG, OB, SS, LP, MMC, IB, PS and DTL carried out experiments. DL performed bioinformatic analyses. IRG, OB, BMS and DTL wrote the paper.

## Abbreviations

ADP: adenosine diphosphate

AMP: adenosine monophosphate

AMPPNP: adenylyl-imidodiphosphate

ATP: adenosine triphosphate

dAMP: deoxyadenosine monophosphate

dATP: deoxyadenosine triphosphate

cryo-EM: cryo-electron microscopy

CTF: contrast transfer function

DNA: deoxyribonucleic acid

HPLC: high pressure liquid chromatography

LB: Luria-Bertani

MWCO: molecular weight cutoff

MST: microscale thermophoresis

PEG: polyethylene glycol

PES: polyether sulphone

PMSF: phenylmethylsulphonyl fluoride

RNR: ribonucleotide reductase

TCEP: tris(2-carboxyethyl)phosphine

TXRF: total-reflection X-ray fluorescence

## Notes

### Competing Interest Statement

The authors have declared no competing interest.

