## Supplementary material for "Bacterial transcriptional repressor NrdR – a flexible multifactorial nucleotide sensor"

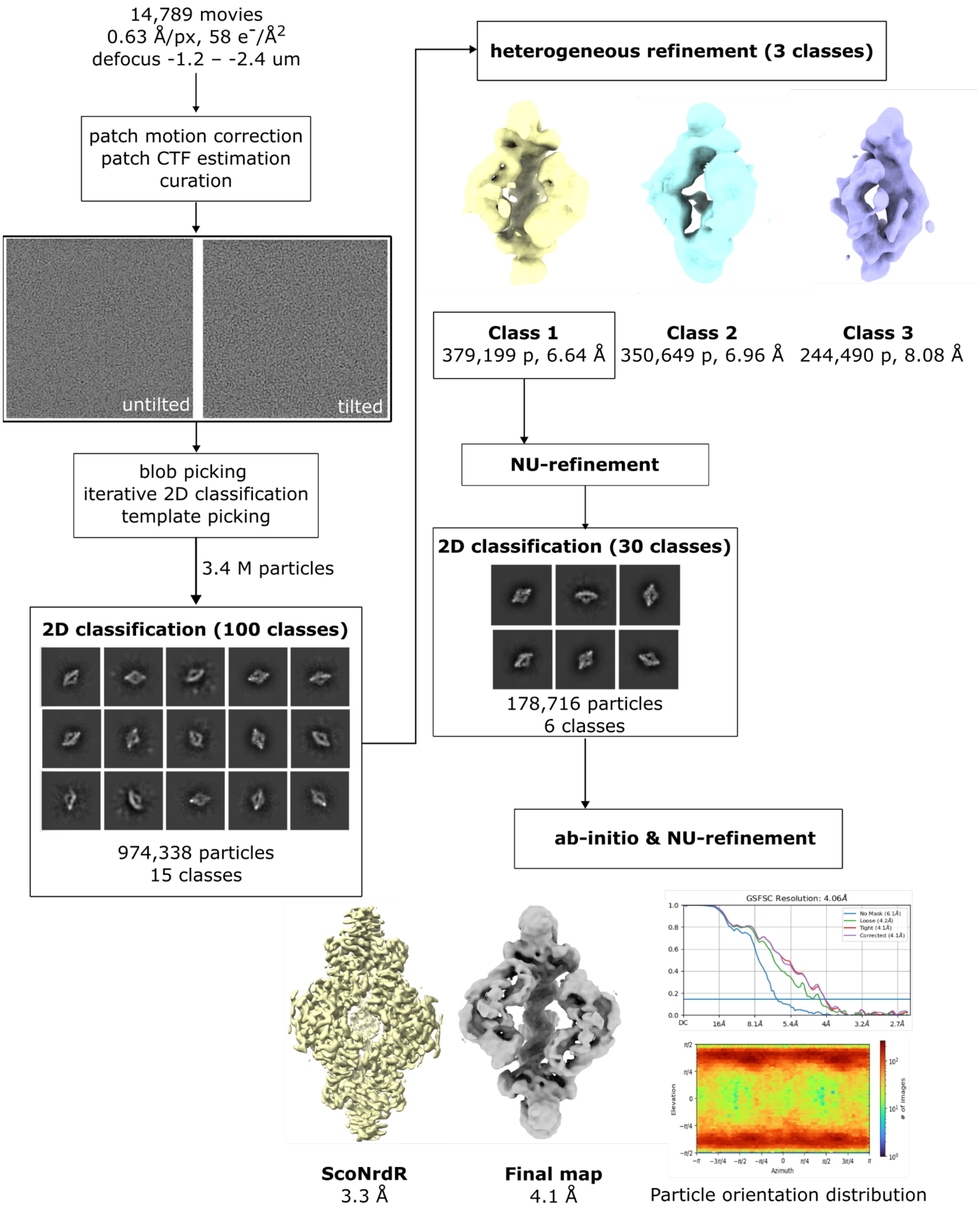

**Supplementary Figure S1. Cryo-EM data processing workflow for the structure of the EcNrdR-ATP-dATP-DNA complex.** The map for the ScoNrdR complex (1) is shown in yellow for comparison.

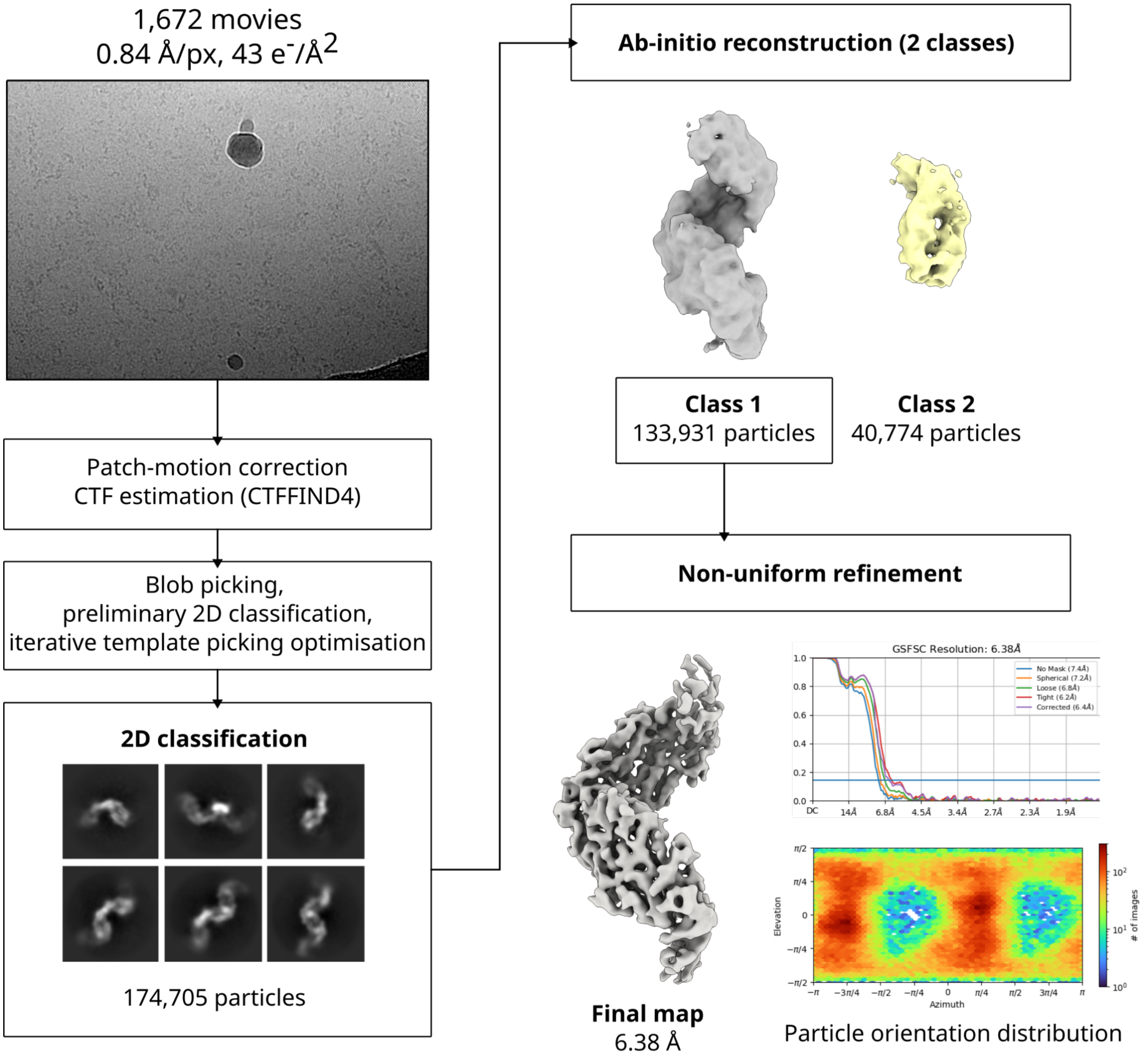

**Supplementary Figure S2. Data acquisition parameters and data processing workflow for the ATP-bound NrdR filament cryo-EM dataset.**

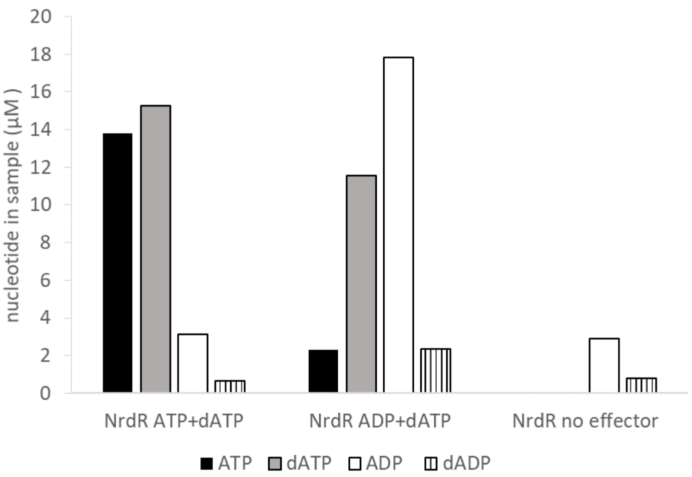

**Supplementary Figure S3. Retained nucleotides in *E. coli* NrdR (35 µM) after addition of effectors and desalting.** 100 µM of as purified-NrdR in Tris-HCl 50mM pH 8.5 at 4 C, 300mM NaCl, 10 mM MgCl2, 0.5 mM TCEP were supplemented with either ATP and dATP or ADP and dATP (1 mM each), incubated at room temperature for 5 minutes and transferred to ice for an additional 20 minutes. A sample without addition of effectors was used as a control. The proteins were desalted from residual nucleotides by applying 100 µl of each sample to a NAP5 column (Cytiva) equilibrated with the same buffer, but without nucleotides. Fractions containing NrdR were collected, and its concentration was determined using Bradford. The samples containing desalted NrdR proteins were boiled for 10 minutes to release the nucleotides bound to it and centrifuged for 10 minutes at 17000 g on a table-top centrifuge. The supernatant was loaded on HPLC (Agilent) using an Agilent ZORBAX RR StableBond (C18, 4.6 x 150 mm, 3.5 µm pore size) equilibrated with buffer A (10% methanol, 50 mM potassium phosphate buffer, pH 7, 10 mM tetrabutylammonium hydroxide). Sample of 10 µL was injected and eluted at 1 mL/min with a gradient of 40%-100% buffer B (30% methanol, 50 mM potassium phosphate buffer, pH 7, 10 mM tetrabutylammonium hydroxide). Compound identification and product quantification based on peak area were performed by external calibration using injected ATP, dATP, ADP, and dADP standards. The correction for phosphate hydrolysis during boiling (10-15% determined experimentally) was taken into account in calculations.

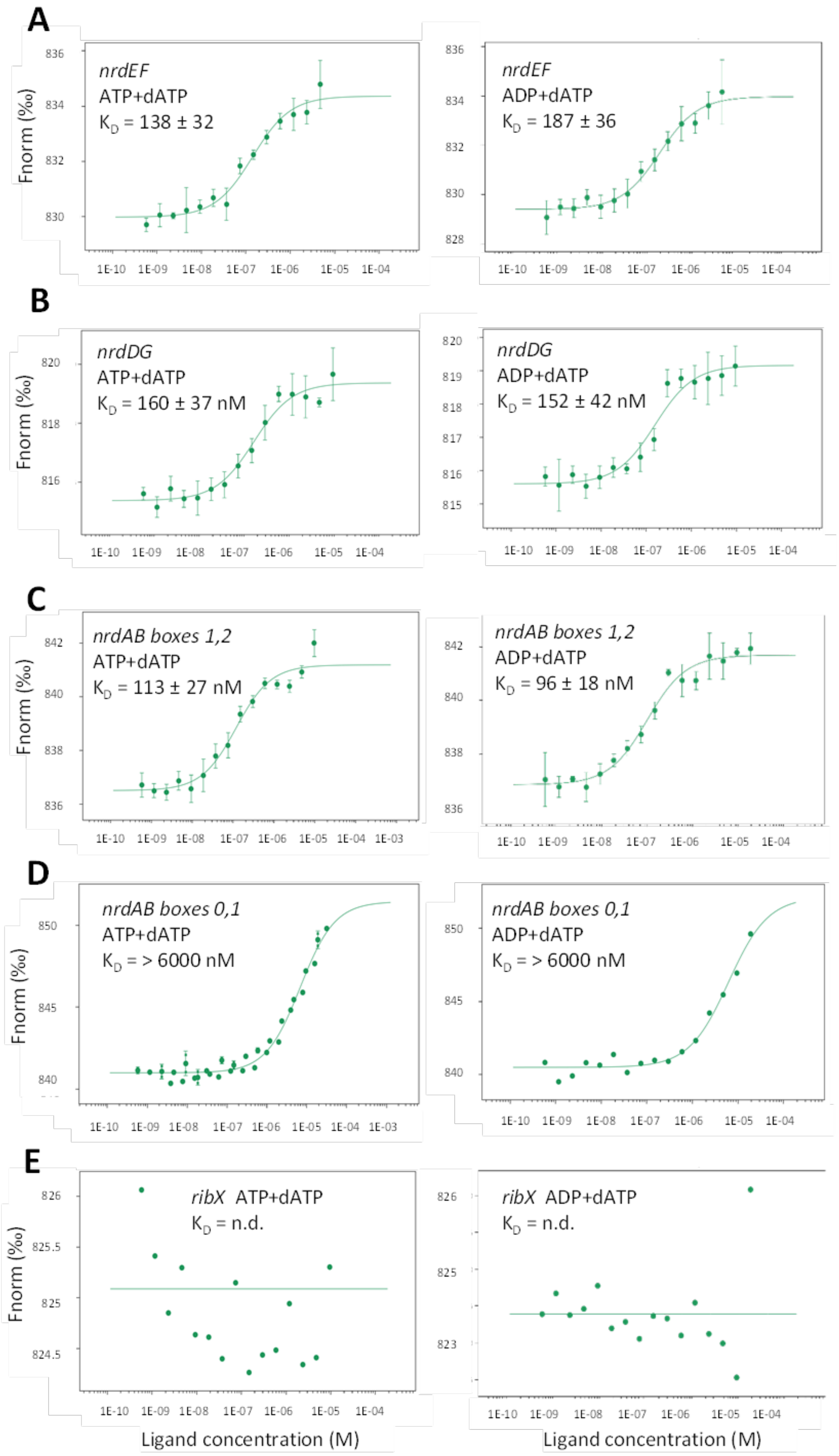

A
nrdEF
ATP+dATP
K_D = 138 ± 32
nrdEF
ADP+dATP
K_D = 187 ± 36
B
nrdDG
ATP+dATP
K_D = 160 ± 37 nM
nrdDG
ADP+dATP
K_D = 152 ± 42 nM
C
nrdAB boxes 1,2
ATP+dATP
K_D = 113 ± 27 nM
nrdAB boxes 1,2
ADP+dATP
K_D = 96 ± 18 nM
D
nrdAB boxes 0,1
ATP+dATP
K_D = > 6000 nM
nrdAB boxes 0,1
ADP+dATP
K_D = > 6000 nM
E
ribX ATP+dATP
K_D = n.d.
ribX ADP+dATP
K_D = n.d.
Fnorm (‰)
Ligand concentration (M)

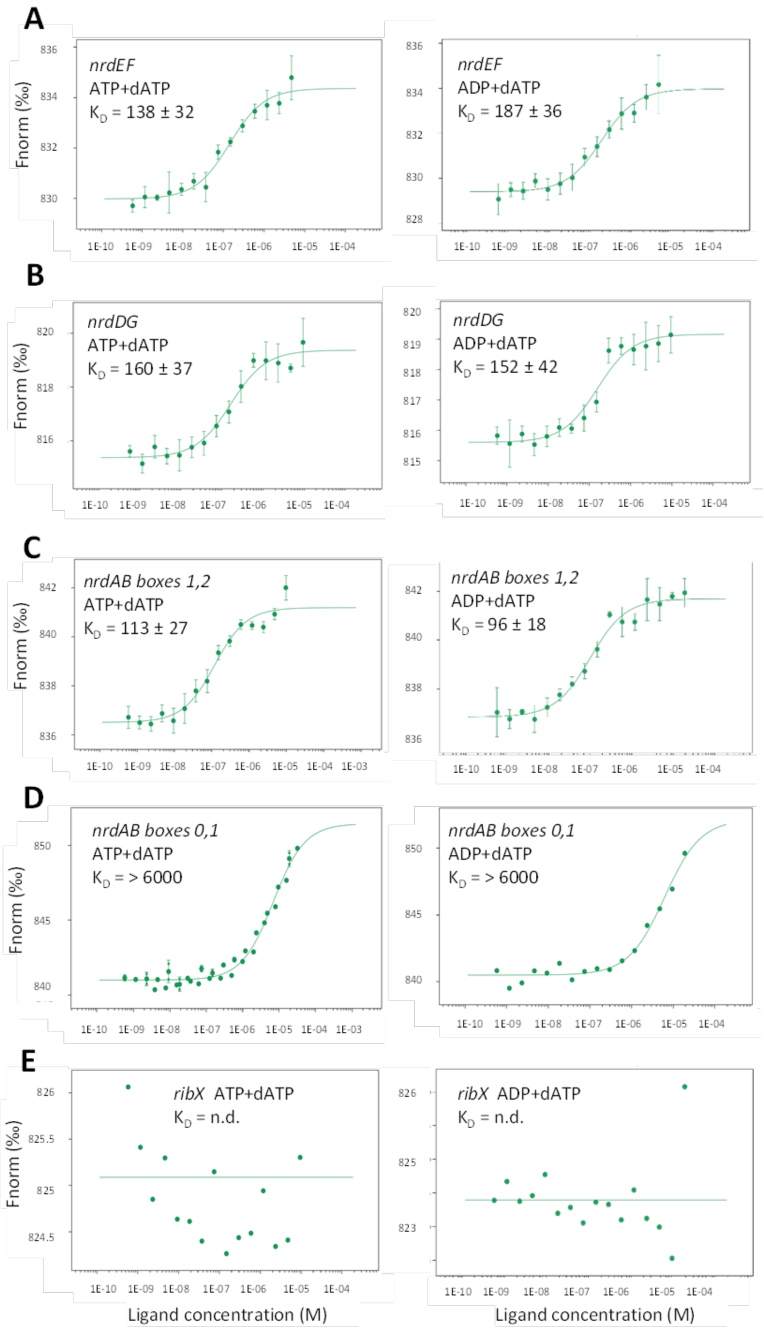

**Supplementary Figure S4. Binding of *E. coli* NrdR simultaneously loaded with dATP and either ATP or ADP to *E. coli* RNR promoters *nrdHIEF* (A), *nrdDG* (B), *nrdAB boxes 1, 2* (C) and alternative *nrdAB boxes 0, 1* (D) and *E. coli ribX* promoter (E) determined by MST.** Plots of the normalized fluorescence Fnorm (‰) from T-Jump and Thermophoresis vs. the concentration of ligand (NrdR) are shown. Lines represent fits of the data points using the KD fit derived from the law of mass action. STDEV derived from at least three experimental repeats. In the case of nrdAB promoter oligo including NrdR boxes 0 and 1 the fits resulted in $K_D$s in the range of 6 μM, but the actual $K_D$ cannot be determined, since the curves do not reach a plateau.

**Supplementary Table S1. Promoter binding constants for effector nucleotide loaded *E. coli* NrdR.**

| **Effector** | ***nrdHIEF* promoter** $K_D$ [nM] | ***nrdDG* promoter** $K_D$ [nM] | ***nrdAB* promoter box 1+2** $K_D$ [nM] | ***nrdAB* promoter box 0+1** $K_D$ [nM] |
|---|---|---|---|---|
| ATP + dATP | 138 ± 32 | 160 ± 37 | 113 ± 27 | >6000 |
| ADP + dATP | 187 ± 36 | 152 ± 42 | 96 ± 18 | > 6000 |
| AMP + dATP | > 5 000 | > 5 000 | > 10 000 | > 10 000 |
| ATP + dADP | n.d. | n.d. | - | n.d. |
| ADP + dADP | 328 | 171 | 128 | > 3600 |
| AMP + dADP | n.d. | n.d. | - | n.d. |
| ATP + dAMP | n.d. | n.d. | - | n.d. |
| ADP + dAMP | n.d. | n.d. | - | n.d. |
| AMP + dAMP | n.d. | n.d. | - | n.d. |
| ATP | n.d. | n.d. | n.d | n.d. |
| ADP | > 100 000 | n.d. | > 5 000 | n.d. |
| AMP | n.d. | n.d. | - | n.d. |
| dATP | > 20 000 | > 6 000 | - | n.d. |
| dADP | n.d. | n.d. | - | n.d. |
| dAMP | n.d. | n.d. | - | n.d. |
| no effector | n.d. | n.d. | n.d | n.d. |
| cAMP | n.d. | - | - | - |
| Cyclic di-AMP | n.d. | n.d. | - | n.d. |
| Cyclic di-AMP + ADP | n.d. | > 100 000 | - | n.d. |

n.d.; not detected; -; not tested

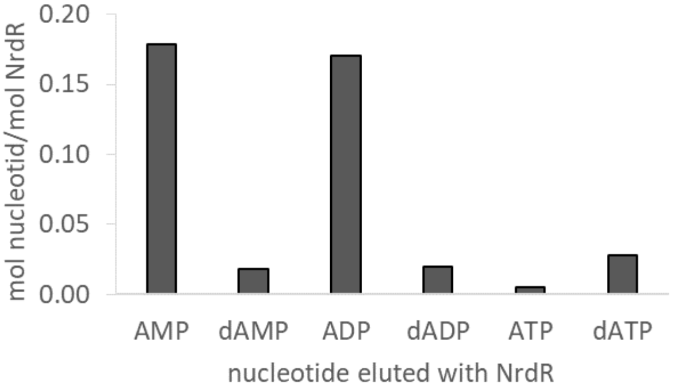

**Supplementary figure S5. Nucleotides eluted with NrdR, analysed using HPLC.** 9 mg/ml (495 µM) recombinant *E. coli* NrdR sample after nickel affinity purification and desalting was boiled for 10 minutes, centrifuged for 10 minutes at 17000 g on a table-top centrifuge, and loaded on HPLC (Agilent) using an Agilent ZORBAX RR StableBond (C18, 4.6 x 150 mm, 3.5 µm pore size) equilibrated with buffer A (10% methanol, 50 mM potassium phosphate buffer, pH 7, 10 mM tetrabutylammonium hydroxide). Sample of 10 µL was injected and eluted at 1 mL/min with a gradient of 40%-100% buffer B (30% methanol, 50 mM potassium phosphate buffer, pH 7, 10 mM tetrabutylammonium hydroxide). Compound identification and product quantification based on peak area were performed by external calibration using injected ATP, dATP, ADP, dADP, AMP and dAMP standards.

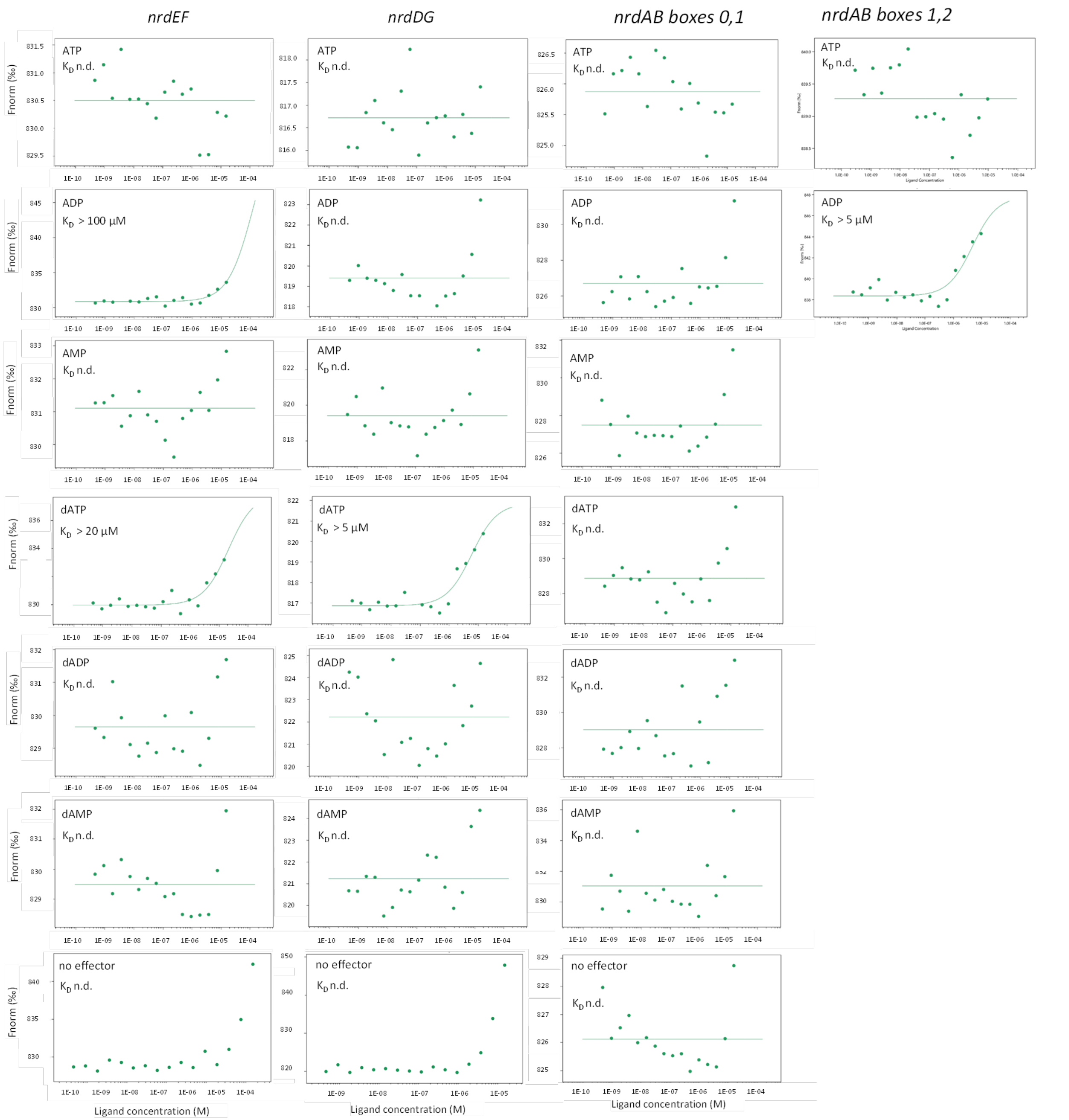

**Supplementary Figure S6. Binding of *E. coli* NrdR loaded with different single adenosine nucleoside phosphates or without any effector (lowest row) to NrdR boxes in *nrdAB*, *nrdHIEF* and *nrdDG* promoters, determined by MST.** Plots of the normalised fluorescence Fnorm (‰) from T-Jump and Thermophoresis vs. the concentration of ligand (NrdR) are shown. Lines represent fits of the data points using the Kd fit derived from the law of mass action. Flat lines were produced in the cases where no fit could be generated by the software and the parameters therefore were fixed as for non-binder ligands. In most cases no binding was detected, and no fit could be generated and in individual cases the fits resulted in $K_D$s in the range of 5 - 100 µM. The actual $K_D$ cannot be determined, since the curves do not reach a plateau. Since these fitted $K_D$s reflect binding affinities lower compared to those of dATP + ATP and dATP + ADP loaded NrdR (see table 1 in the main text), we believe that they are not physiologically relevant and reflect non-specific binding. n.d. not determined.

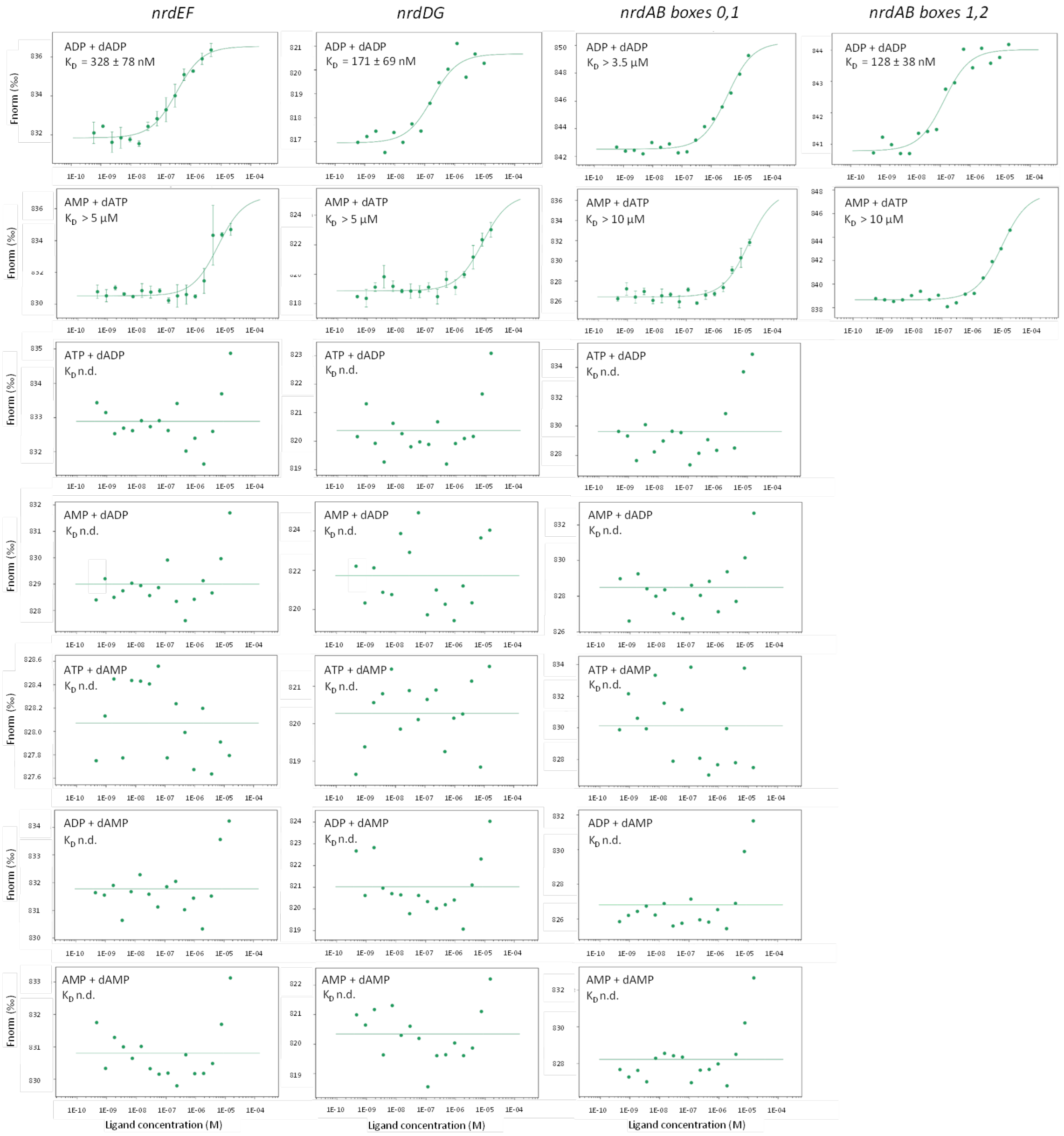

**Supplementary Figure S7. Binding of *E. coli* NrdR loaded with all possible combinations of adenosine nucleoside phosphates to NrdR boxes in *nrdAB*, *nrdHIEF* and *nrdDG* promoters, determined by MST.** Plots of the normalised fluorescence Fnorm (‰) from T-Jump and Thermophoresis vs. the concentration of ligand (NrdR) are shown. Lines represent fits of the data points using the Kd fit derived from the law of mass action. Flat lines were produced in the cases where no fit could be generated by the software and the parameters therefore were fixed as for non-binder ligands. In some cases the fits resulted in $K_D$s in the range of 3.5 - 10 μM, but the actual $K_D$ cannot be determined, since the curves do not reach a plateau. Since these fitted $K_D$s reflect binding affinities lower compared to those of dATP + ATP and dATP + ADP loaded NrdR (see table 1 in the main text), we believe that they are not physiologically relevant and reflect non-specific binding. In many cases no binding was detected, and $K_D$ couldn't be fitted and therefore they are marked n.d. not determined.

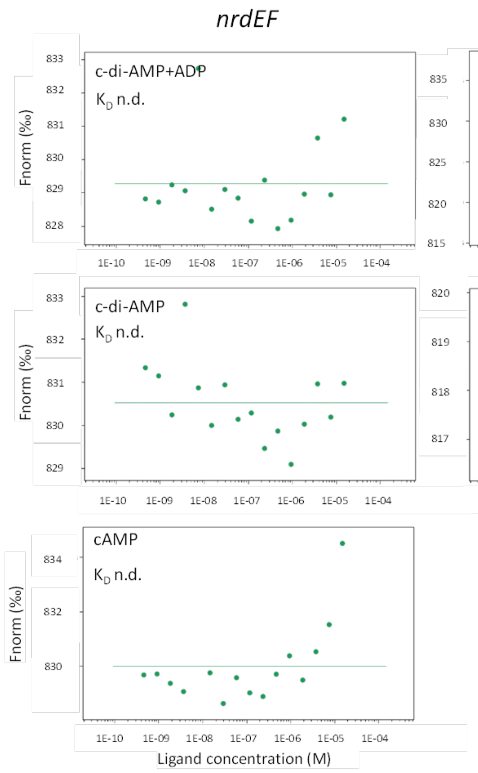

**Supplementary Figure S8. Binding of *E. coli* NrdR loaded with c-di-AMP, a combination of c-di-AMP and ADP and cAMP to NrdR boxes in *nrdAB*, *nrdHIEF* and *nrdDG* promoters, determined by MST.** Plots of the normalised fluorescence Fnorm (‰) from T-Jump and Thermophoresis vs. the concentration of ligand (NrdR) are shown. Lines represent fits of the data points using the Kd fit derived from the law of mass action. Flat lines were produced in the cases where no fit could be generated by the software and the parameters therefore were fixed as for non-binder ligands. In the case of c-di-AMP- and ADP-loaded NrdR and the nrdDG promoter, the fit resulted in $K_D$s higher than 100 µM and most likely reflecting non-specific binding. The actual $K_D$ cannot be determined, since the curve doesn't reach a plateau. Binding of NrdR loaded with cyclic AMP to DNA was assayed only with *nrdHIEF* promoter region. n.d. not determined.

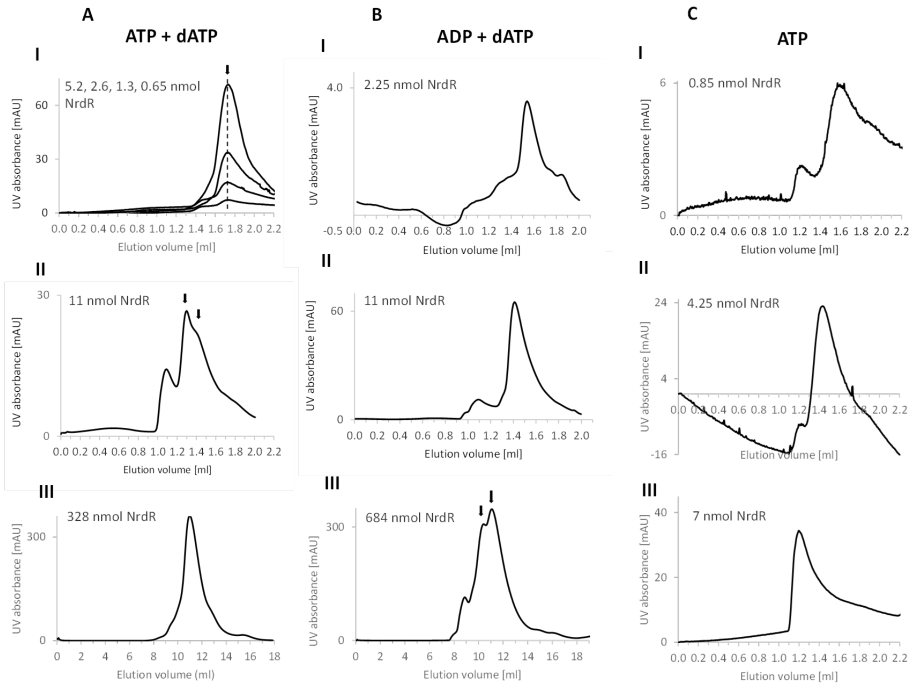

IV

| panel | Mw (Kda) | # of subunits |
|---|---|---|
| I | 83 | 4.5 |
| II | 156, 260 | 8.3, 13.9 |
| III | 278 | 15.2 |

IV

| panel | Mw (Kda) | # of subunits |
|---|---|---|
| I | 76 | 4.1 |
| II | 148 | 7.9 |
| III | 267, 367 | 14.6, 20 |

IV

| panel | Mw (Kda) | # of subunits |
|---|---|---|
| I | 214 | 12 |
| II | 430 | 24 |
| III | n.d. (v.v.) | > 33 |

**Supplementary figure S9. Size exclusion chromatography of NrdR with different effectors at varying protein quantities.** Representative chromatograms are shown. **A.** NrdR supplemented with ATP and dATP. **AI**: 95 µl of 55, 27, 14 and µM NrdR samples were loaded on a Superdex 200 PC 3.2/30 column connected to Shimadzu HPLC system. **AII**: 25 µl of 440 µM NrdR were loaded on Superdex 200 PC 3.2/30 column connected to an ÄKTA prime system. **AIII**: 0.6 ml of 547 µM NrdR was loaded to Superdex 200 gl 10/300 increase column (for purification prior to crystallisation) **B:** NrdR supplemented with ADP and dATP. **BI, BII**: 25 µl of 90 and 440 µM NrdR respectively were loaded on Superdex 200 PC 3.2/30 column connected to Äkta prime system. **BIII:** 1 ml of 684 µM NrdR was loaded on Superdex 200 gl 10/300 increase column (for purification prior to crystallisation) **C:** NrdR supplemented with ATP CI, CII, CIII: 50 µl of 17, 85 and 140 µM NrdR samples were loaded on a Superdex 200 PC 3.2/30 column connected to Shimadzu HPLC system equilibrated with buffer containing low MgCl2 and ATP. All NrdR samples were incubated with respective effector nucleotides before SEC and nucleotides were included in SEC buffer in all cases. See Materials and methods for further SEC details. Arrows indicate the peaks for which molecular weight was calculated. **AIV, BIV, CIV:** summary of eluted NrdR complexes. Molecular weight of eluted NrdR complexes calculated based on standards for each column. Number of subunits was calculated based on molecular weight of NrdR of 18 280 Da.

**Supplementary Table S2. Data collection and refinement statistics**

| | ADP/dATP | AMPPNP/dATP | ATP/dATP-SeMet |
|---|---|---|---|
| **Data collection** | | | |
| Beamline | DLS I04 | MAX IV BioMAX | MAX IV BioMAX |
| Wavelength (Å) | 0.9795 | 0.9763 | 0.9791 |
| Space group | $P6_222$ | $P2_12_12_1$ | $P2_12_12_1$ |
| Cell dimensions | | | |
| *a*, *b*, *c* (Å) | 147.1 147.1 85.8 | 84.8 129.8 143.9 | 70.8 73.8 138.0 |
| *α*, *β*, *γ* (°) | 90, 90, 120 | 90, 90, 90 | 90, 90, 90 |
| Anisotropic resolution limits | 3.53, 3.53, 2.27 | 2.56, 2.33, 2.67 | – |
| Resolution (Å) [a] | 127.4-2.44 (2.70-2.44) | 71.92-2.33 (2.42-2.33) | 65.10-3.10 (3.37-3.10) |
| Number of unique reflections | 10312 (517) | 51826 (2591) | 10661 (534) |
| Multiplicity | 39.1 (41.2) | 13.7 (13.6) | 13.1 (12.3) |
| $R_{merge (I)}$ [a] | 0.310 (2.635) | 0.204 (2.451) | 0.215 (2.082) |
| $R_{pim}$ [a] | 0.050 (0.413) | 0.057 (0.687) | 0.062 (0.606) |
| Mean I/σ(I) [a] | 10.6 (1.6) | 10.0 (1.5) | 6.77 (0.40) |
| $CC_{1/2}$ [a] | 0.999 (0.912) | 0.996 (0.427) | 0.997 (0.117) |
| Spherical completeness (%)[a] | 49.6 (9.9) | 75.0 (14.5) | 78.29 (9.93) |
| Ellipsoidal completeness (%)[a] | 95.1 (84.7) | 99.4 (55.8) | – |
| **Refinement** | | | |
| Resolution (Å) | 127.4-2.44 (2.67-2.44) | 96.37-2.33 (2.48-2.33) | 65.11-3.11 (3.22-3.11) |
| No. reflections | 10312 (419) | 51862 (1037) | 10639 |
| $R_{free}$ / $R_{work}$ | 0.283 / 0.215 (0.324 / 0.272) | 0.265 / 0.228 (0.447 / 0.343) | 0.287 / 0.213 (0.297 / 0.388) |
| No. atoms | | | |
| protein | 2404 | 9656 | 4878 |
| nucleotides | 116 | | |
| $Zn^{2+}$ ions | 2 | 8 | 4 |
| water | 161 | 115 | |
| Average B factors (Å$^2$) | | | |
| protein | 49.5 | 66.4 | 106.5 |
| nucleotides | | | |
| $Zn^{2+}$ ions | | 176.9 | |

| | | | |
|---|---|---|---|
| water | 29.0 | 62 .5 | 60.5 |
| R.m.s. deviations | | | |
| bond lengths (Å) | 0.007 | 0.013 | 0.006 |
| bond angles (°) | 0.9 | 1.6 | 1.3 |
| Ramachandran plot | | | |
| favoured (%) | 96.9 | 0.979 | 0.866 |
| outliers (%) | 1.4 | 0 | 0.34 |
| All-atom clash score (percentile) | 4.6 (99th) | | |

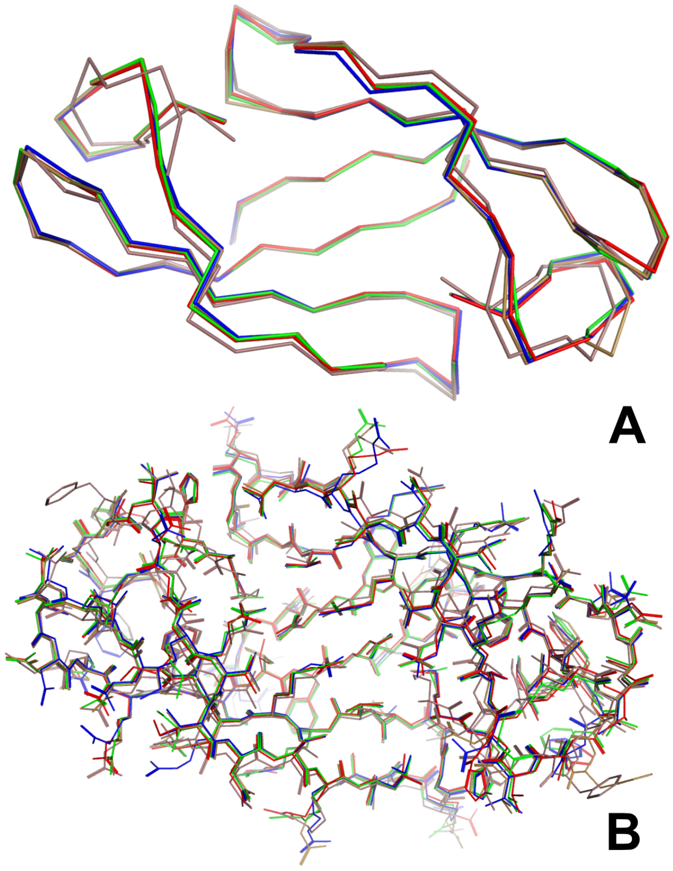

**Supplementary figure S10. Superposition of all Zn-ribbon domain pairs from the crystal structures of the AMPPNP-dATP-bound and ADP-dATP-bound forms of EcoNrdR**. A) backbone trace; B) all atoms. The four pairs from the AMPPNP-dATP-bound form are coloured red, green, blue and beige respectively, while the single pair from the ADP-dATP-bound form is coloured light purple. The deviations between the structures are almost exclusively in the side chains on the surface. A small deviation between the ADP-dATP form and the other pairs is seen in a loop on the top left / bottom right of the dimer, but this is most likely due to crystal packing effects. The structures support the observation that the Zn-ribbon pairs behave as rigid bodies.

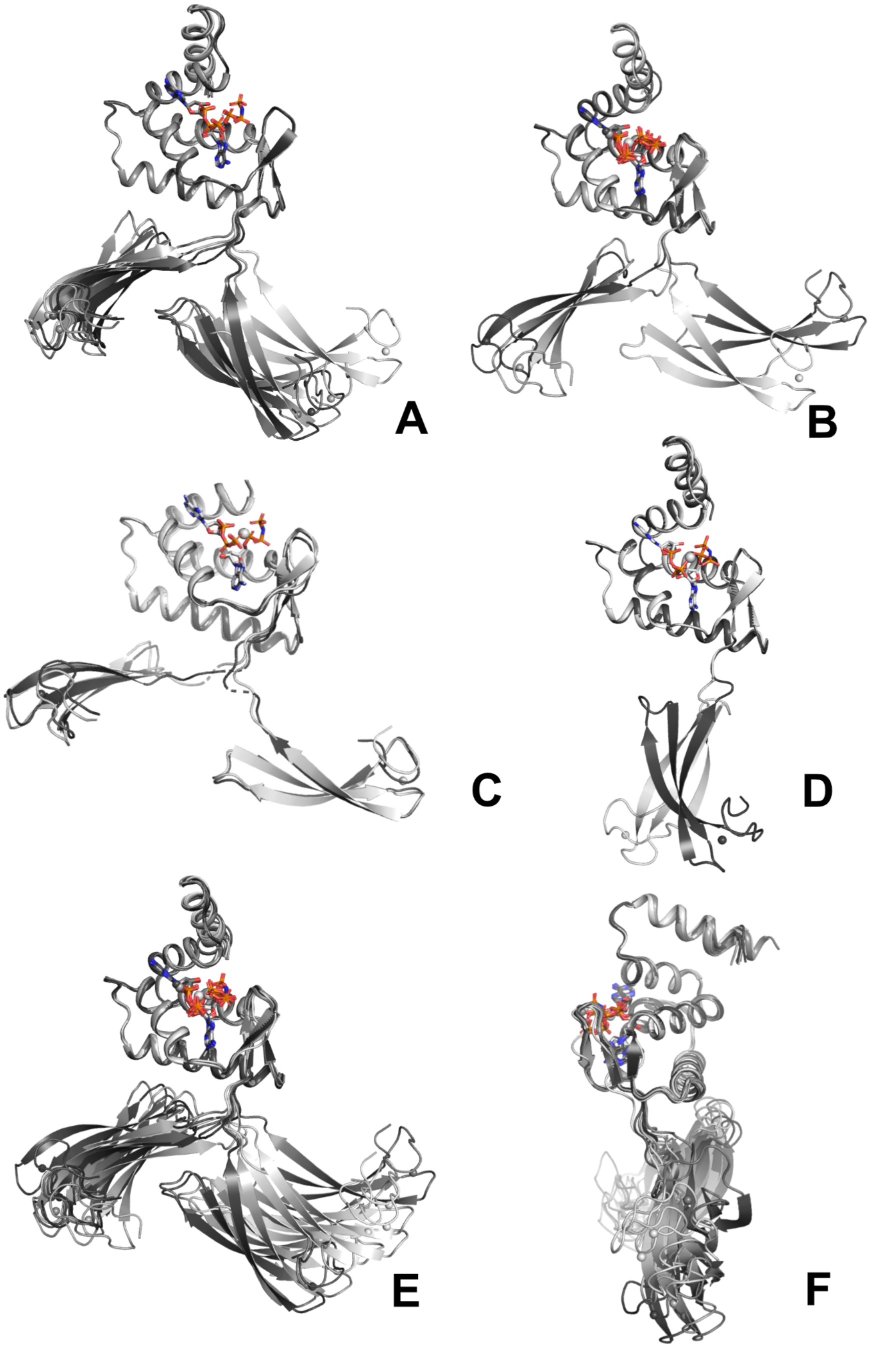

**Supplementary figure S11. The angles between ATP-cone and Zn-ribbon domains form clusters.** A) The eight polypeptide chains in the two independent tetramers of EcoNrdR-AMPPNP-dATP in the asymmetric unit, superimposed on the ATP-cone domains; B) The four chains of the single tetramer in the asymmetric unit of SeMet-EcoNrdR-ATP-dATP; C) Two chains from the EcoNrdR-ATP filament; D; The two independent chains of the EcoNrdR-ATP-dATP-DNA complex (refined with C2 symmetry); E) and F) Two orthogonal views of all 12 chains of structures except the DNA complex. This is in effect a superposition of panels A, B and C. The nucleotides in all structures are shown as sticks.

**Supplementary Table S3. Angles (in degrees) between ATP-cone and Zn-ribbon domains in the crystal- and cryo-EM structures of EcoNrdR reported in this work, as well as the previously published structures of ScoNrdR.** The angle reported is the one subtended by the CA atoms of residues 81 and 98 (which define the ends of the helix in the ATP-cone closest to the Zn-ribbon domain) and the Zn atom of the Zn-ribbon (Figure 4C). For ScoNrdR the equivalent helix begins and ends at residues 78 and 95. Average values and standard deviations for the clusters formed by chains of type A and type B are given in the last four columns. For the ATP filament, only the angles for the central two tetramers are given, as the fit to the map is best in that region.

| | A | B | C | D | E | F | G | H | average cluster A | average cluster B |
|---|---|---|---|---|---|---|---|---|---|---|
| **EcoNrdR** | | | | | | | | | | |
| **AMPPNP-dATP tetramer** | 131.9 | 39.8 | 107.8 | 45.5 | 107.8 | 41.3 | 108.8 | 41.9 | 114.1 ± 10.3 | 42.1 ± 2.6 |
| **ATP-dATP tetramer** | 128.6 | 49.2 | 151.2 | 40.0 | | | | | 139.9 ± 11.3 | 44.6 ± 4.6 |
| **ATP-dATP-DNA** | 91.1 | 63.3 | 92.3 | 64.3 | | | | | 92.0 ± 0.3 | 63.8 ± 0.5 |
| **ADP-dATP tetramer*** | 109.8 | 34.9 | | | | | | | 109.8 | 34.9 |
| **ATP filament** | 136.6 | 35.4 | 137.6 | 34.2 | 138.3 | 35.2 | 139.9 | 36.3 | 138.1 ± 1.2 | 35.3 ± 0.9 |
| **ScoNrdR** | | | | | | | | | | |
| **ATP-dATP octamer** | 88.0 | 75.1 | | | | | | | 88.0 | 75.1 |
| **ATP-dATP-DNA** | 85.9 | 81.8 | 86.0 | 77.2 | | | | | 86.0 ± 0.1 | 79.5 ± 2.3 |
| **ATP dodecamer** | 131.1 | 68.8 | | | | | | | 131.1 | 68.8 |

* Chain B forms the compact tetramer in the ADP-dATP complex, chain A the infinite filaments.

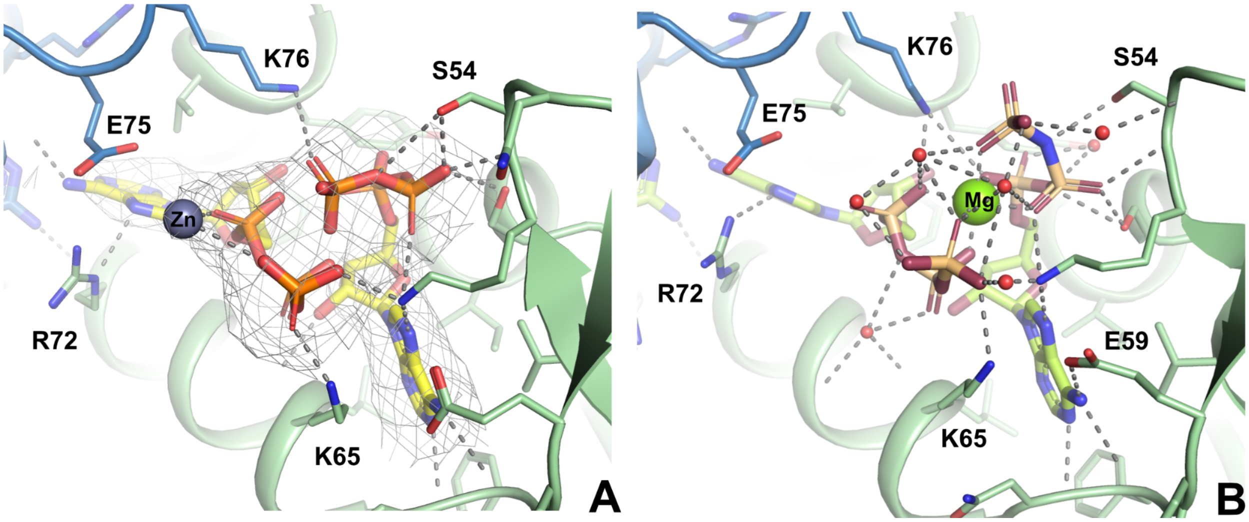

**Supplementary Figure S12. A) Electron density map for the EcoNrdR-dATP-ATP complex and binding of ATP** (right) **and dATP** (left)**. B) Comparison of AMPPNP** (right) **and ATP** (left) **binding for comparison.**

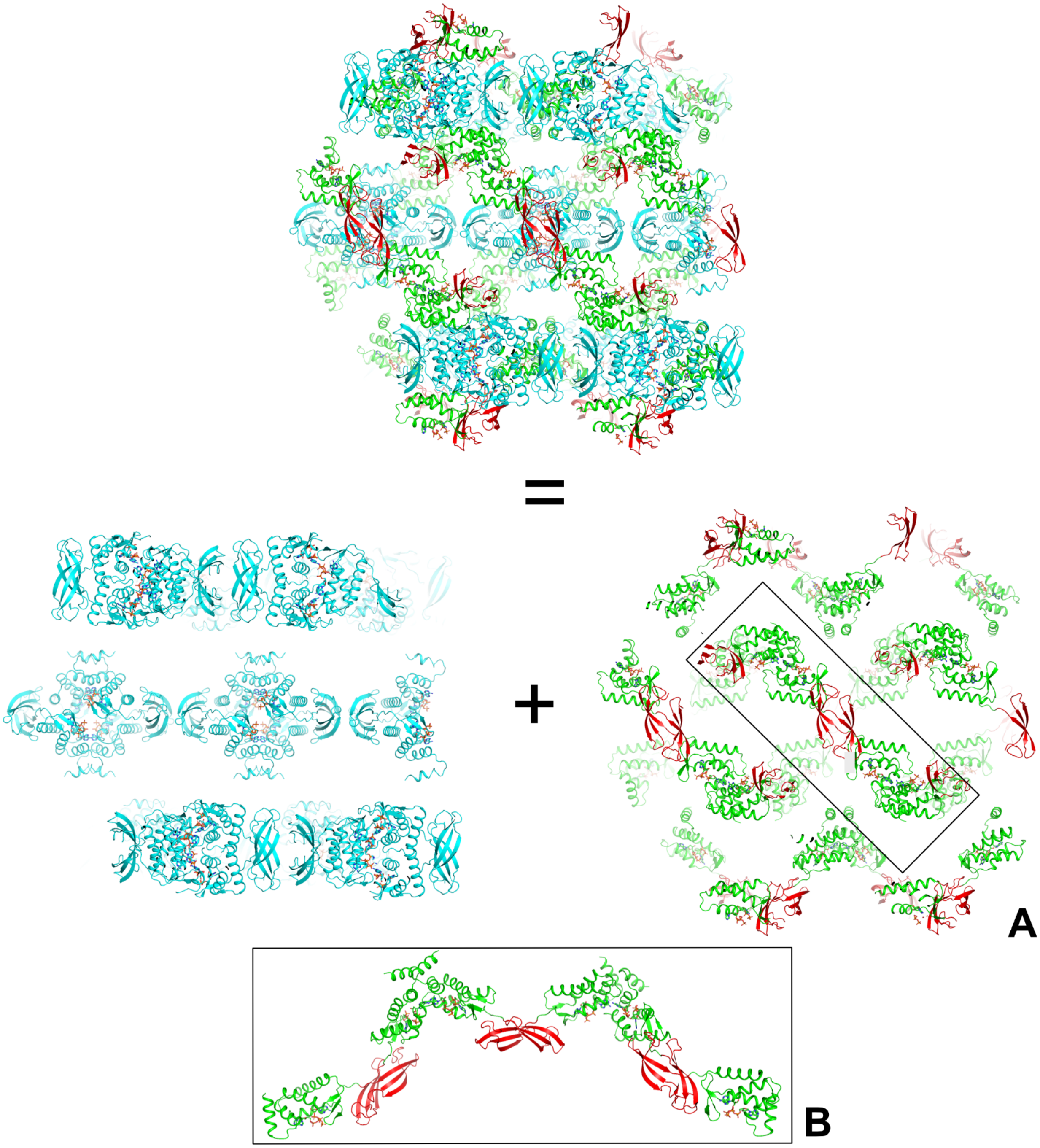

**Supplementary Figure S13. Crystal packing in the ADP/dATP-bound form of EcoNrdR.** A) The crystal packing can be decomposed into compact tetramers (blue) decorated with infinite chains of monomers linked by alternating interactions of pairs of ATP-cone domains (light blue) and Zn-ribbon domains (red). B) Zoom in on part of the infinite chain.

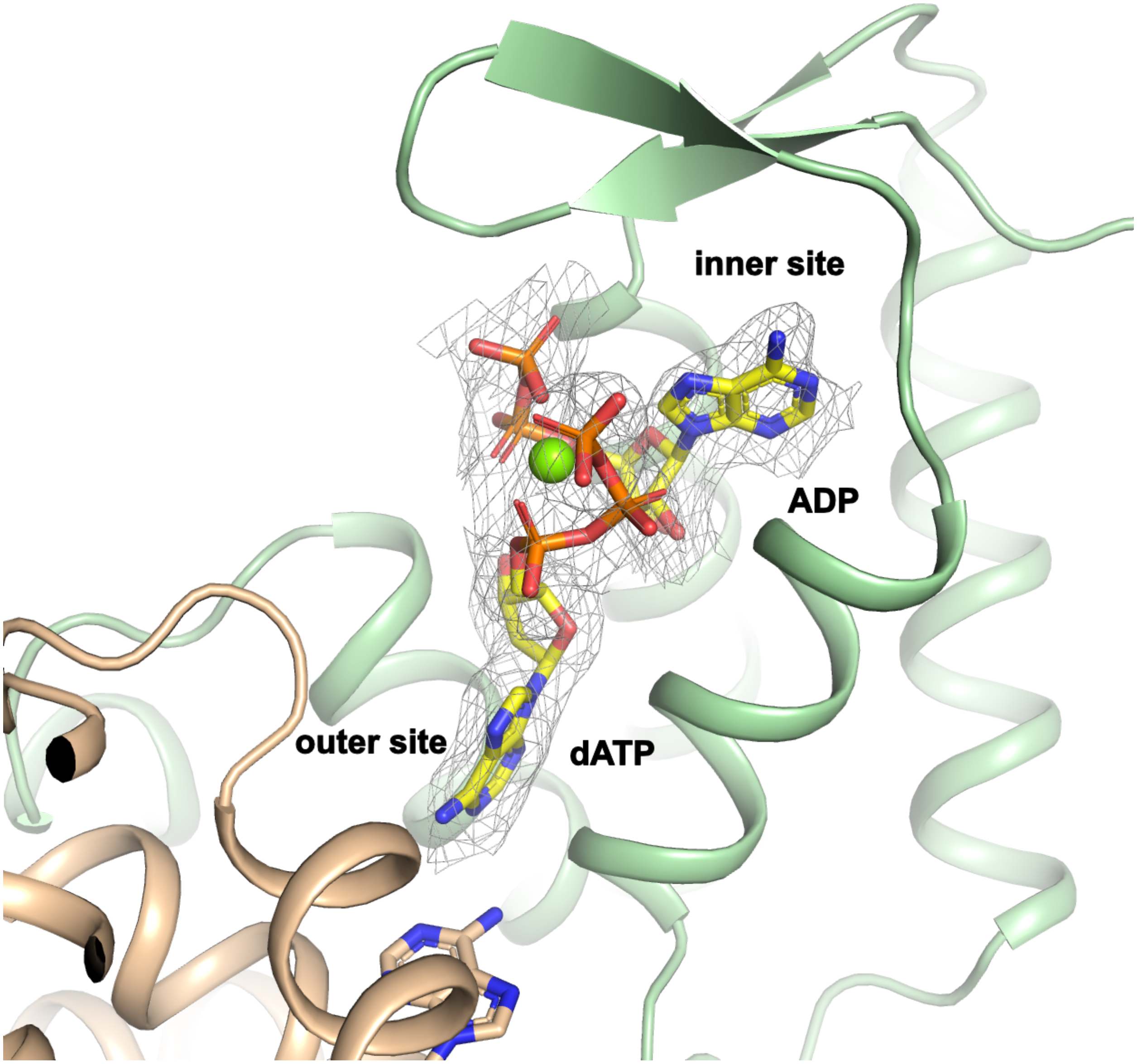

**Supplementary figure S14. Nucleotides in ADP/dATP structure.** Overall view of one ATP-cone (green) showing the inner and outer sites binding ADP and dATP respectively. The second ATP-cone in the dimer is shown in beige. The $Mg^{2+}$ ion is drawn as a green sphere. 2m|Fo|-D|Fc| electron density for the nucleotides contoured at 1.1 σ is shown as a grey mesh.

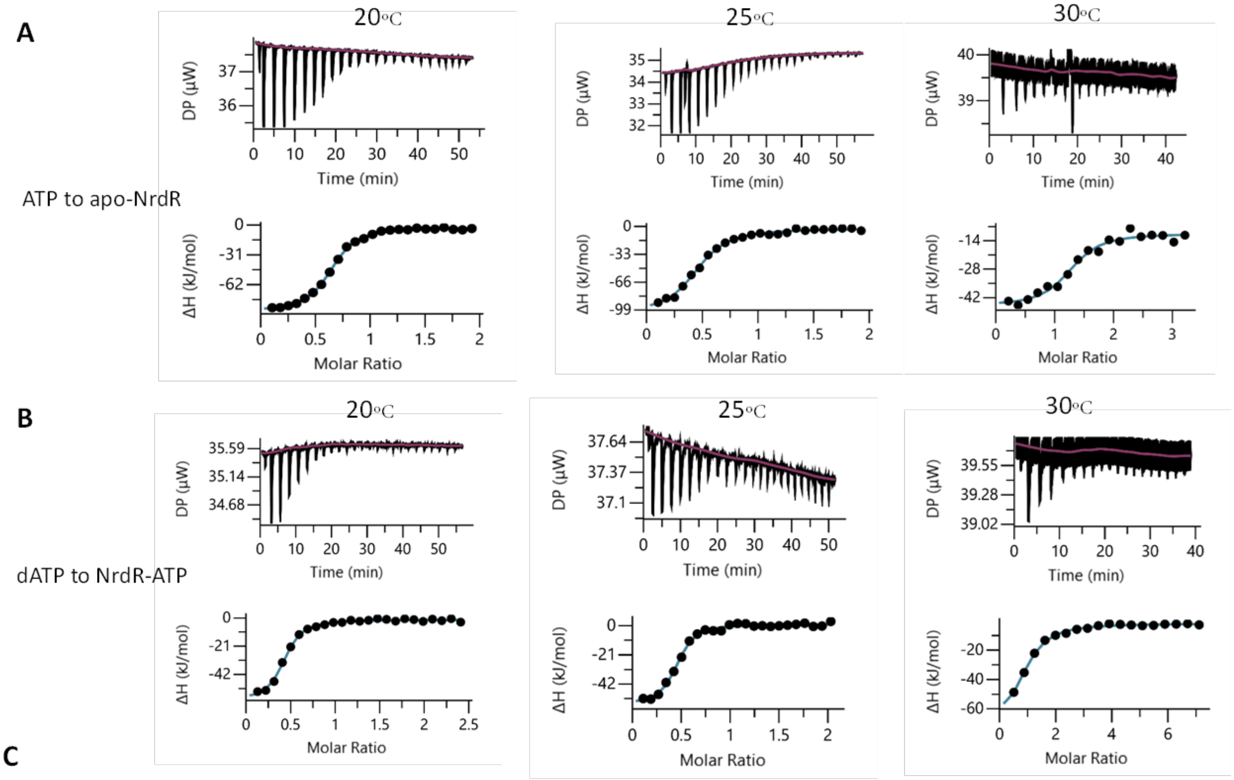

C

| Temperature (°C) | protein | Ligand | N | Kd (μM) | ΔH (kJ/mol) | ΔG (kJ/mol) | -TΔS (kJ/mol) |
|---|---|---|---|---|---|---|---|
| 20 | apo-NrdR | ATP | 0.48 ± 0.13 | 0.377 ± 143 | -88 ±2.5 | -36 ± 1.1 | 51 ± 1.6 |
| 20 | NrdR-ATP | dATP | 0.426 | 0.387 | -57.9 | -36 | 21.9 |
| 25 | apo-NrdR | ATP | 0.445 | 0.987 | -101 | -34.3 | 66.8 |
| 25 | NrdR-ATP | dATP | 0.395 | 0.413 | -60.2 | -36.5 | 23.8 |
| 30 | apo-NrdR | ATP | 1.23 | 0.337 | -35.7 | -37.6 | -1.9 |
| 30 | NrdR-ATP | dATP | 0.809 | 1.1 | -75.2 | -34.4 | 40.6 |

**Supplementary Figure S15. ITC analyses of ligand binding to *E. coli* NrdR at 20 °C, 25 °C and 30 °C.** Representative ITC thermograms obtained by titration of ATP to apo- NrdR (A) and of dATP to NrdR loaded with ATP (B). Isothermal calorimetric enthalpy changes (upper panels) and resulting binding isotherms (lower panels) are shown. (C) Thermodynamic parameters of ligand binding to NrdR. Binding isotherms were fitted using a one-set-of-sites binding model. Values for ATP titration of apo-NrdR at 20 °C reported as the mean ±SD of three titrations. Other titrations are based on a single binding experiment for each ligand at each temperature. All titrations were performed as described in Methods.

**List of supplementary movies:**

**Supplementary movie S1. Conformational differences between the EcoNrdR and ScoNrdR complexes with dATP-ATP-DNA** (1). The morph and movie were created using ChimeraX v1.7.1 (2). The movie starts with the ScoNrdD structure, morphs to the EcoNrdR structure and back again.

**Supplementary movie S2. Conformational changes in EcoNrdR upon DNA binding, starting from the conformation in the dATP-AMPPNP tetramer.** The movie shows the adjustment necessary for EcoNrdR to attain a conformation where it can bind two NrdR boxes simultaneously, assuming a hypothetical starting state in which one pair of Zn ribbons is bound to one NrdR box. The DNA is in its final, highly bent form and is included as reference.

**Supplementary movie S3. Conformational changes in EcoNrdR upon DNA binding, starting from the conformation in the ADP-dATP tetramer.** The movie shows the adjustment necessary for EcoNrdR to attain a conformation where it can bind two NrdR boxes simultaneously, assuming a hypothetical starting state in which only one pair of Zn ribbons is bound to one NrdR box. The DNA is in its final, highly bent form andis  included as reference.

**Supplementary movie S4. Conformational changes between the EcoNrdR-AMPPNP-dATP tetramer and the tetramer observed in the EcoNrdR-ATP filament.** The movie starts with the AMPPNP-bound state and ends with the ATP-bound state. The ATP-cones rotate relative to each other while the angle between ATP-cone and Zn-ribbons remains similar.

**Supplementary Movie S5. Hypothetical conformational change between a state where one Zn-ribbon pair of EcoNrdR in the dATP-ATP-bound state is bound to one NrdR box in undistorted B-DNA and the final state with highly curved DNA and both Zn-ribbon pairs bound to NrdR boxes.** The B-DNA was generated from the sequence of the coding strand of the DNA in the ScoNrdR complex using ChimeraX v1.7.1 (2).

Supplementary Movies S2-S5 were made using PyMOL v2.5.8 (3). Intermediate states are interpolated and do not necessarily represent true intermediates, especially for the DNA in Supplementary Movie S5.
